# Spatial Transcriptomics and Single-Nucleus RNA Sequencing Reveal rAAV2- and rAAV9-Specific Transduction Signatures in the Mouse Liver

**DOI:** 10.1101/2025.03.13.643011

**Authors:** Bettina Amberg, Fabian Köchl, Nadine Kumpesa, Megana Prasad, Filip Bochner, Mar Hernández-Obiols, Michael B. Otteneder, Lucas Stalder, Frances Shaffo, Ali Nowrouzi, David Markusic, Hélène Haegel, Rebecca Xicluna, Marc Sultan, Björn Jacobsen, Sven Rottenberg, Alberto Valdeolivas, Petra C. Schwalie, Kerstin Hahn

**Author notes:** These authors jointly supervised this work (Alberto Valdeolivas, Petra C. Schwalie, Kerstin Hahn). Corresponding author Correspondence: Kerstin Hahn, Roche Pharma Research and Early Development, Roche Innovation Center Basel, Grenzacherstrasse 124, Basel CH-4070, Switzerland.

## Abstract

The liver is a primary target for recombinant adeno-associated viral (rAAV) vectors, yet the influence of serotype, sex, and liver zonation on transduction and transcriptomic changes remain incompletely understood. This proof-of-concept study employs spatial transcriptomics alongside single-nucleus RNA sequencing to map the spatial distribution and impacts of rAAV2- and rAAV9-CMV-EGFP vectors in male and female mouse livers. Spatial transcriptomics provided precise transgene mapping and highlighted that rAAV vectors deregulate hepatocellular lipid metabolism, the circadian clock, and the immune/stress response with sex specific differences. Lipid metabolism genes (*Elovl3*, *Chka*, *Irs2*, *Ppard*), were deregulated independent of zonation, serotype, and sex, while *Srebf1*, *Tlcd4*, *Cpt2*, and *Acot1* exhibited sex-specific patterns. Circadian clock modulators (*Dbp*, *Tef*, *Arntl*, *Nfil3*, *Nr1d1/Nr1d2*) were altered independent of zonation. The study found sex-specific downregulation of immune and stress-response genes and pathways, including *Gadd45g* and hypoxia pathways. TGF-β and EGFR pathways were upregulated sex-independently. Spatial transcriptomics further enabled examination of transgene and rAAV entry factor co-expression, identifying known and novel factors like *Rpsa*, *Dpp4*, *Sdc1*, and solute carrier proteins highlighting its role in supporting targeted screening. Our findings demonstrate spatial transcriptomics as a powerful tool in gene therapy research and reveal novel rAAV vector effects on liver biology.

**Graphical Abstract:** 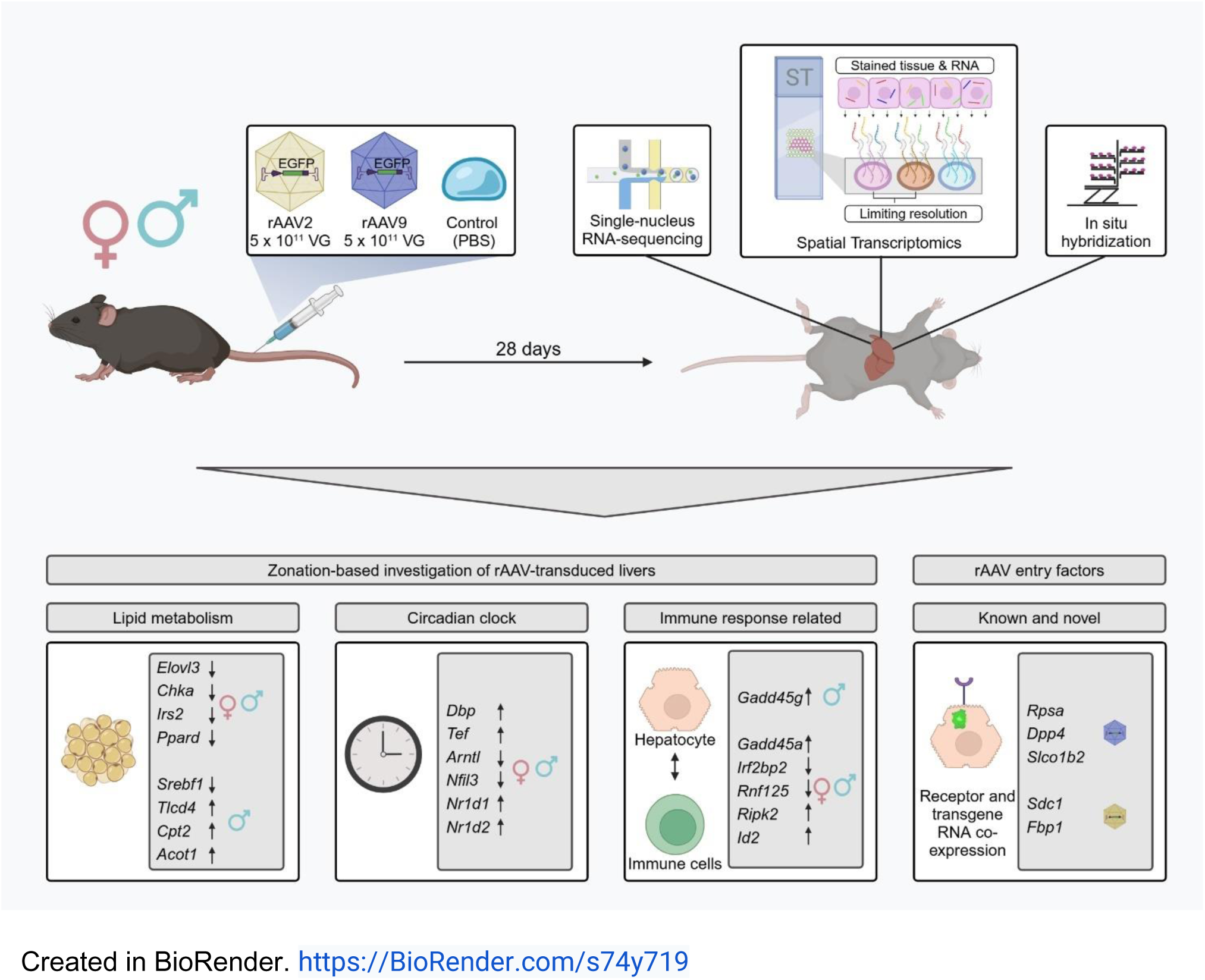

## Introduction

The liver is a primary target or off-target organ for gene therapy using recombinant adeno-associated viral (rAAV) vectors.^1^ The liver is composed of lobes, which are further divided into repeating structural units called lobules. Each lobule is subdivided into zones—periportal, intermediate, and pericentral— each with distinct metabolic functions. These zones exhibit a gradient of enzyme expression and metabolic activities, allowing different regions of the lobule to specialize in various biochemical processes such as gluconeogenesis, glycolysis, and detoxification.^2^ In mice, the spatial distribution of rAAV gene therapy vectors is related to liver zonation and is influenced by both serotype and sex. Whereas rAAV2-based vectors preferentially transduce periportal regions, rAAV9 favors pericentral zones.^3^ Overall, liver transduction is generally higher in male mice.^4,5^

These sex- and serotype-specific differences may stem from androgen-regulated pathways that influence rAAV entry, trafficking, genome processing, or immune modulation.^5^ Additionally, differences in liver zonation-specific metabolism and AAV receptor/co-receptor expression may further contribute to these variations. The differences in liver transduction lead to varying transgene expression levels in male and female mice, potentially causing distinct downstream effects and potential toxicities. In this regard, methods that link transgene expression with genome-wide transcriptomic changes *in situ* could help better characterize AAV transduction and its associated transcriptomic alterations.

Here, we apply 10x Genomics Visium spatial transcriptomics (ST) and single-nucleus RNA sequencing (snRNA-seq) to comprehensively map rAAV2- and rAAV9-mediated transduction in the mouse liver. Unlike classical methods mapping rAAV biodistribution as immunohistochemistry (IHC) or *in situ* hybridization (ISH), ST uniquely enables spatial mapping of transgene expression to tissue structures and analysis of associated transcriptomic signatures. Furthermore, compared to panel-based, multiplex approaches in rAAV research, such as USeqFISH^6^ or STARmap PLUS^7^, which analyze 50 or 1000 genes, 10x Genomics Visium ST enables unbiased, whole-transcriptome profiling. We reasoned that this combination offers novel opportunities for hypothesis generation, specifically for understanding both the cellular responses to rAAV transduction and factors influencing rAAV entry.

In our proof-of-concept study, we applied Visium ST to delineate differentially expressed genes, pathway activity, and receptor expression associated with rAAV transduction in the liver of male and female mice transduced with rAAV2 or rAAV9 encoding enhanced green fluorescent protein (EGFP) under a cytomegalovirus (CMV) promoter (rAAV2-CMV-EGFP and rAAV9-CMV-EGFP). To cross-validate and resolve findings at the single-cell level, we utilized snRNA-seq analyses that were carried out using the same liver samples.

Our study establishes 10x Genomics Visium as a powerful platform for spatially mapping transgene expression in the mouse liver. We demonstrate that rAAV2- and rAAV9-mediated transduction induces broad, zonation- and sex-independent modulation of the circadian clock in liver regions with high EGFP transgene expression. Additionally, we uncover deregulation of lipid metabolism, replication stress responses, and innate antiviral immunity, with distinct sex-specific patterns. Through a comprehensive ST analysis, we also validate known rAAV2 and rAAV9 co-receptors and propose putative novel entry factors. These findings provide a proof of concept for using spatially resolved transcriptomics to elucidate molecular signatures relevant to gene therapy, offering insights into vector tropism, host responses, and potential targets for optimizing AAV-based liver-directed therapies.

## Results

### Spatial transcriptomics maps rAAV biodistribution

We hypothesized that the 10x Genomics Visium ST platform represents a novel method for mapping rAAV vectors to tissue-specific hallmarks and assessing corresponding transcriptomic signatures at the whole-genome level. To validate our approach, we analyzed liver samples from male and female mice transduced with either 5 x 10^11^ vector genomes (VG) rAAV2- or rAAV9-CMV-EGFP, along with control samples. For these serotypes distinct spatial distribution patterns have been described.^3^ We complemented our ST analysis with snRNA-seq and confirmed transgene expression using ISH (Figure 1A, Methods).

**Figure 1:**
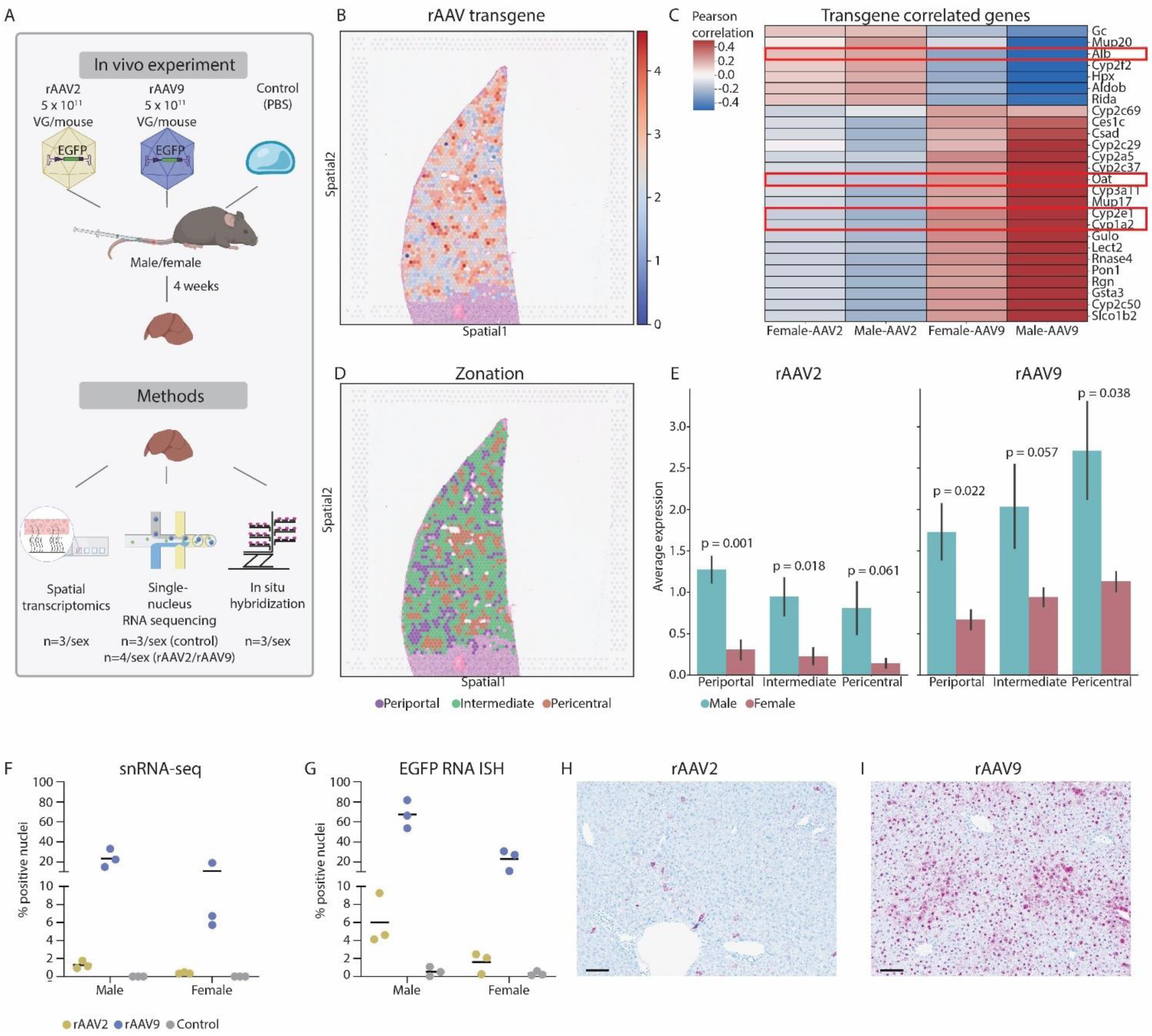
Spatial transcriptomics deciphers rAAV liver biodistribution. **(A)** Experimental setup. Created in BioRender. https://BioRender.com/q78t856 **(B)** Spatial mapping of the transgene expression in the liver of a rAAV9-treated male mouse. **(C)** Pearson’s correlation between transgene abundance and gene expression across rAAV9- and rAAV2-treated male and female liver samples. Known periportal and pericentral zonation markers are highlighted. **(D)** Spatial mapping of the predicted zones in the liver of a male, rAAV9-treated mouse. **(E)** Average transgene expression for rAAV9 and rAAV2 per zone in male and female mice. Black bars represent the standard deviation per condition. Statistical analysis to compare males and females was performed using Welch’s t-test (Methods), P values are indicated. **(F)** Percentage of transgene positive nuclei in the single-nucleus RNA sequencing data (snRNA-seq) for male and female rAAV2-, rAAV9-treated and control animals. Each dot represents one individual with the mean represented by horizontal lines. Only animals with corresponding spatial transcriptomics, *In situ* hybridization and EGFP immunohistochemistry data are displayed. **(G)** *In situ* hybridization (ISH) using an EGFP antisense probe maps the vector in liver tissue. Graph shows quantification of ISH as percent cells showing cytoplasmatic ISH signals, as well as both cytoplasmic and nuclear ISH signals, corresponding to RNA expression. Cells showing exclusively nuclear ISH signals, suggestive of rAAV DNA, were not included. Each dot represents one individual with the mean represented by horizontal lines. **(H)** EGFP antisense ISH of the liver of an rAAV2-treated male animal. Scale bar 100 μm. **(I)** EGFP antisense ISH of the liver of an rAAV9-treated male animal. Scale bar 100 μm.

ST detected the transgene across the whole liver section (Figure 1B). We employed Pearson correlation analysis to identify genes that are positively associated with the rAAV transgene, revealing numerous liver zonation markers (Figure 1C). To examine the zonal distribution of the transgene, the different liver zones were assigned using conserved markers (Figure 1D, Figure S1A-F, Methods).^2,8–10^ Zone classifications were assessed by a pathologist based on morphological features and further validated by using known zonation markers (Figure S1 A-D). Moreover, we applied deconvolution, a widely used computational technique for estimating cell type proportions in ST data, to map periportal and pericentral hepatocytes from the snRNA-seq data to the tissue.^11^ Periportal and pericentral hepatocytes aligned with periportal and pericentral zones, respectively (Figure S1E-F, Methods). Animals treated with rAAV9 showed higher pericentral transgene expression, while rAAV2 exhibited the highest transgene expression in the periportal zone (Figure 1E). Additionally, animals treated with rAAV9 had higher transgene expression than those treated with rAAV2, and males demonstrated higher transgene expression than females for both rAAV2 and rAAV9 (Figure 1E). These findings are consistent with previously reported results.^3^ snRNA-seq confirmed the sex- and serotype-specific patterns (Figure 1F). The analysis of transgene-expressing cells revealed that around 50-60% are hepatocytes (Figure S1G-J and Table S1). ISH and IHC confirmed the differences between sexes and serotypes (Figure 1G-I and Figure S1K). These findings were further in line with In Vivo Imaging System (IVIS) data generated in an independent experiment (Figure S1L-O).

Hence, we demonstrate the ability of 10x Genomics Visium ST to spatially map the transgene expression and confirm the distinct sex- and serotype-specific patterns of rAAV vector biodistribution and transgene expression in the liver, with rAAV9 favoring the pericentral zone and rAAV2 the periportal zone.

### ST and snRNA-seq enable zonation-based investigation of rAAV-transduced livers

ST provides a comprehensive analysis of both nuclear and cytoplasmic RNA signatures, offering superior transgene capture while mitigating the ambient RNA contamination often observed in liver snRNA-seq data.^12^ Given these advantages, we focused on ST data to dissect liver zonation-, serotype-, and sex-specific patterns in male and female mice treated with rAAV2- or rAAV9-CMV-EGFP. To identify differentially expressed genes (DEGs), we compared transcriptomic profiles across periportal, intermediate and pericentral zones of treated and control animals within the same sex and assessed common DEG signatures. To align these findings with snRNA-seq data, we performed a parallel analysis of periportal and pericentral hepatocytes (Figure S2A, Methods).^9,10^ Our analysis identified numerous DEGs associated with lipid metabolism, the circadian clock, and immune response modulation, with representative findings shown in Figure 2A. To further explore underlying regulatory mechanisms, we applied the decoupleR package to infer transcription factor and pathway activity based on observed gene expression patterns and prior knowledge on regulons and signaling pathways (Figure 3 and Figure S3, Methods).^13^ Additionally, Gene set enrichment analysis (GSEA) was applied on the Gene Ontology Biological processes and the Kyoto Encyclopedia of Genes and Genomes (KEGG) gene sets to identify enriched functional pathways (Figure 4 and Figure S4, Methods).^14,15^ To enhance the robustness of our findings, we prioritized genes exhibiting consistent trends across both ST and snRNA-seq datasets. This integrated approach allowed us to highlight the most biologically relevant changes, which are explored in detail in the following sections.

**Figure 2:**
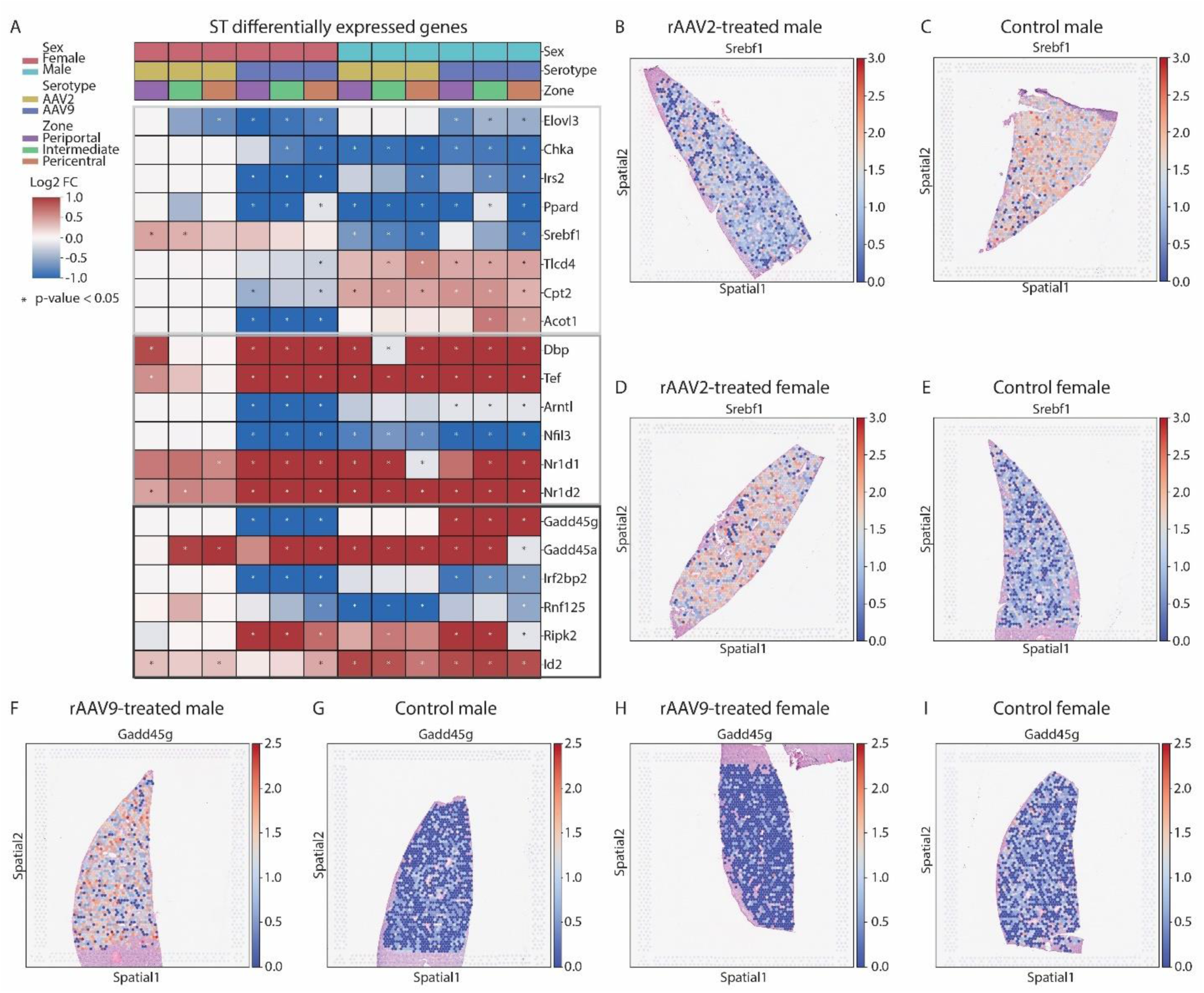
Spatial transcriptomics reveals differential gene expression in rAAV2- and rAAV9-treated mice livers per zone. **(A)** Differential gene expression in treated versus control animals, categorized by liver zone and sex in the spatial transcriptomics (ST) data. Displayed is a selection of genes related to lipid metabolism (light grey square), circadian rhythm (medium gray square), and immune modulation (dark grey square). P values were calculated using the Wald statistical test (Methods). **(B-E)** Spatial mapping of *Srebf1* gene expression across the liver of **(B)** a male rAAV2-treated animal, **(C)** a male control animal, **(D)** a female rAAV2-treated animal and **(E)** a female control animal. **(F-I)** Spatial mapping of *Gadd45g* gene expression across the liver of **(F)** a male rAAV9-treated animal, **(G)** a male control animal, **(H)** a female rAAV9-treated animal and **(I)** a female control animal.

**Figure 3:**
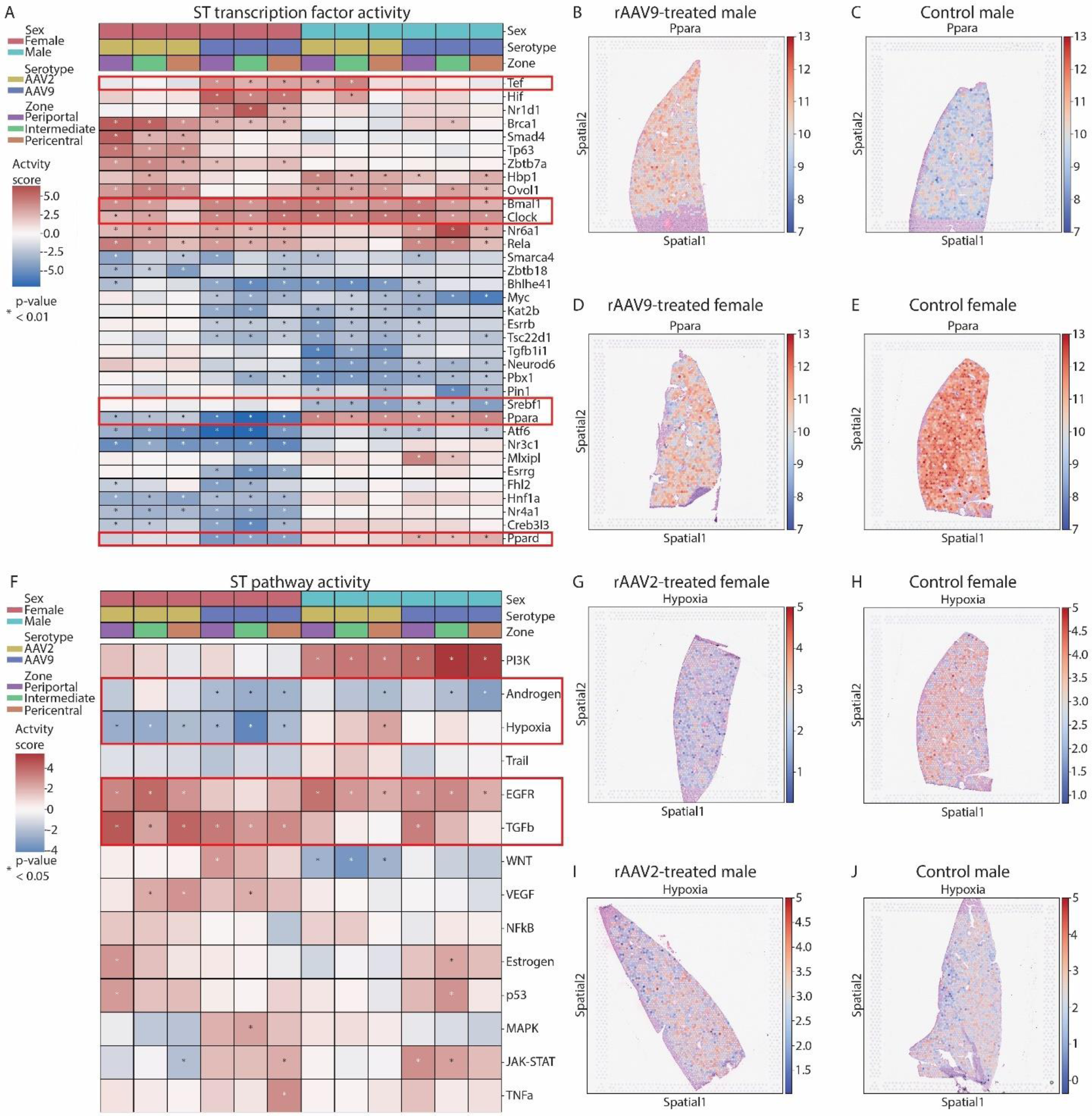
Spatial transcriptomics reveals transcription factor and pathways activity changes in mice livers per zone. **(A)** Predicted transcription factor activity per zone in the spatial transcriptomics (ST) data. P values were calculated using the Wald statistical test (Methods). Genes discussed in the text are highlighted with a red square. **(B-E)** Spatial mapping of the predicted Ppara transcription factor activity across the liver of **(B)** a male rAAV9-treated animal, **(C)** a male control animal, **(D)** a female rAAV9-treated animal and **(E)** a female control animal. **(F)** Differential pathway activity computed on pseudo-bulkRNA-seq generated from the zones in the ST data. P values were calculated using the Wald statistical test (Methods). Pathways discussed in the text are highlighted with a red square. **(G-J)** Spatial mapping of the predicted hypoxia pathway activity across the liver of **(G)** a female rAAV2-treated animal, **(H)** a female control animal, **(I)** a male rAAV2-treated animal and **(J)** a male control animal.

**Figure 4:**
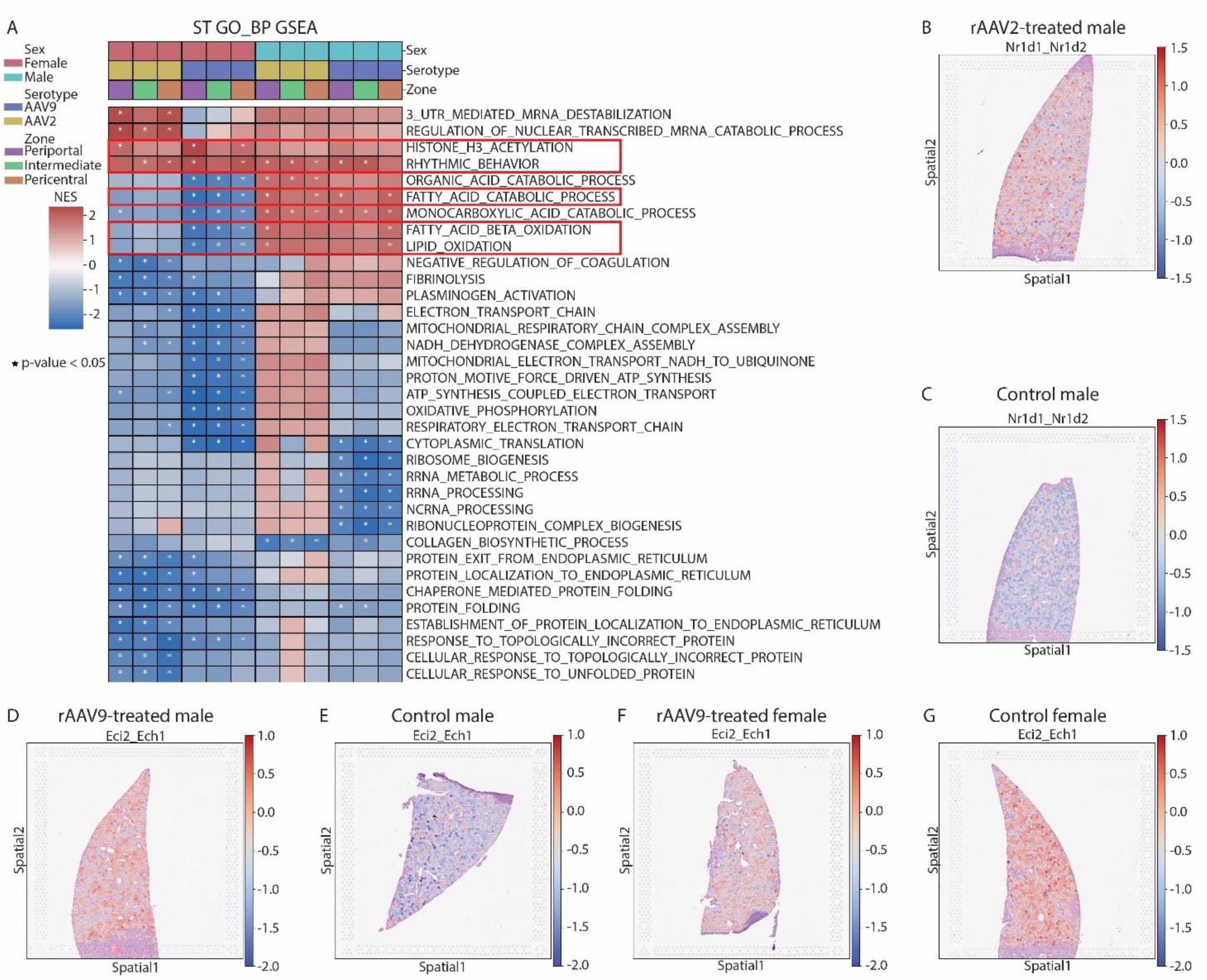
Spatial transcriptomics reveals changes in rhythmic processes and lipid metabolism. **(A)** Gene set enrichment analysis (GSEA) using the GO biological process (GO_BP) database in the spatial transcriptomics (ST) data. NES = normalized enrichment score. ‘REGULATION_OF_NUCLEAR_TRANSCRIBED_MRNA_CATABOLIC_ PROCESS _DEADENYLATION_DEPENDENT_DECAY‘ is shortened for display purposes. P values were calculated using the Wald statistical test (Methods). Gene sets discussed in the text are highlighted with a red square. **(B-C)** Combined score of the two leading genes *Nr1d1* and *Nr1d2* for the ‘RHYTHMIC_BEHAVIOUR‘ process across the liver of **(B)** a male rAAV2-treated and **(C)** a male control animal. **(D-G)** Combined score of the two leading genes *Eci2* and *Ech1* for the ‘FATTY_ACID_CATABOLIC_PROCESS‘ and ‘FATTY_ACID_BETA_OXIDATION‘ across the liver of **(D)** a male rAAV9-treated animal, **(E)** a male control animal, **(F)** a female rAAV9-treated animal and **(G)** a female control animal.

#### rAAV2- and rAAV9-CMV-EGFP is associated with zonation-independent changes in the lipid metabolism

ST revealed alterations in lipid metabolism-associated genes, displaying partially sex-specific but overall serotype- and zonation-independent patterns.

Assessment of DEGs indicated overall a downregulation of *Elovl3*, a long-chain fatty acid elongase,^16^ that was most pronounced in rAAV9-treated males and females. *Chka,* involved in synthesis of lipid membrane components,^17^ and *Irs2*, which plays a role in fatty acid synthesis,^18^ were downregulated in rAAV2- and rAAV9-treated males and rAAV9-treated females, exhibiting varying intensities of deregulation across different conditions/zones (Figure 2A). *Ppard,* which regulates fatty acid beta oxidation and induces antiviral effects,^19,20^ was significantly downregulated in rAAV2- and rAAV9- treated males and rAAV9-treated females (Figure 2A).

Transcription factor activity estimates showed Ppard significantly upregulated in rAAV9-treated males and downregulated in rAAV9-treated females suggesting compensatory regulation of upstream components and indicating sex-specific pathways modulating Ppard activity (Figure 3A-E). Similarly, transcription factor activity estimates predicted Ppara, a regulator of lipid metabolism with antiviral effects,^20^ to be significantly upregulated in rAAV2- and rAAV9-treated females and downregulated in males (Figure 3A). *Srebf1* a regulator of fatty acid and cholesterol synthesis^21,22^ was overall downregulated in rAAV2- and rAAV9-treated males in differential expression analysis and transcription factor activity estimates (Figure 2A-E and Figure 3A).

In addition, we identified sex-specific DEG patterns for *Tlcd4* and *Cpt2*, which are involved in lipid metabolism and fatty acid oxidation, respectively.^23–25^ Overall, there was an upregulation of these genes in rAAV2- and rAAV9-treated males, with *Cpt2* showing a significant increase (Figure 2A).

*Acot1*, a regulator of fatty acid oxidation,^26,27^ was upregulated in rAAV9-treated males and downregulated in females (Figure 2A).

Pathway activity scores showed a trend toward downregulation of the androgen pathway, which is closely linked to lipid metabolism, in rAAV2- and rAAV9-treated males and rAAV9-treated females (Figure 3F).

In line with DEG and transcription factor activity prediction, GSEA revealed sex-specific trends including an overall upregulation of fatty acid catabolic processes as well as fatty acid and lipid oxidation in the males and downregulation in the females (Figure 4A, D-G). Additionally, GSEA using the KEGG database revealed an upregulation of arachidonic acid metabolism in rAAV2- and rAAV9-treated males and a downregulation of PPAR signaling in rAAV2- and rAAV9-treated females (Figure S4B). Of note, only females showed downregulation of processes related to electron transport and mitochondrial processes as well as translation and ribosome biogenesis, both linked to energy metabolism (Figure 4A).

In snRNA-seq data, all of the DEGs exhibited the highest or high expression in hepatocytes (Figure S2B) and overall, similar up and/or downregulation patterns were observed for most of the genes (Figure S2A), transcription factor activity estimates (Figure S3A), pathway activity assessments (Figure S3B) and GSEA (Figure S4A, C).

These changes may be linked to rAAV transduction and/or transgene expression, potentially modulating the AAV life cycle, transgene expression levels, and host immune responses in a partially sex-specific manner.

#### rAAV2- and rAAV9-CMV-EGFP modulate the circadian clock across liver zones in male and female mice

Our ST data further revealed several rAAV-related circadian clock-associated signatures, that were overall zonation- and sex-independent and found for both rAAV2 and rAAV9.

DEG analysis identified an overall zonation-independent upregulation of the circadian clock regulators *Dbp* and *Tef^28^* in rAAV9-treated mice and rAAV2-treated males (Figure 2A), which was significant in periportal and pericentral zones. Conversely, the circadian rhythm regulator *Arntl* (encoding Bmal1) and *Nfil3*, a circadian rhythm inhibitor showed a general trend of downregulation, whereas *Nr1d1* and *Nr1d2* were upregulated across liver zones (Figure 2A).^29,30^ Transcription factor activity predictions suggested an overall upregulation of Bmal1 and Clock activity (Figure 3A). These findings indicate an upregulation of genes upstream of Bmal1, coupled with a compensatory downregulation of the *Arntl* transcription itself. In line with the observed changes in the DEG, GSEA predicted an upregulation of rhythmic behavior in both male and female rAAV2- and rAAV9-treated animals (Figure 4A-C), which was consistent with the upregulation of histone H3 acetylation, a process regulated by CLOCK and BMAL1,^30^ in all treated conditions (Figure 4A).

Overall, the snRNA-seq data showed similar trends, and additionally revealed a significant downregulation of *Dbp* and *Tef* for rAAV-2-treated females (Figure S2A).

#### rAAV2 and rAAV9-CMV-EGFP alters immune response-related genes with zonation-independent and partially sex-specific patterns

ST data provided further evidence for rAAV2- and rAAV9-mediated alterations in hepatocyte-driven immune modulation and DNA replication-associated genes.

DEG analysis identified in rAAV9-treated males a significant upregulation of *Gadd45g* (Figure 2A, F-I), which is associated with replication stress and immune response.^31^ In contrast, a significant downregulation of *Gadd45g* was observed in females.

Additionally, *Gadd45a*, involved in immune regulation, DNA repair, and stress responses, was overall upregulated in the ST data (Figure 2A). *Irf2bp2*, a regulator of macrophage function^32,33^ and also expressed in hepatocytes,^8^ was significantly downregulated in rAAV9-treated animals. *Rnf125*, a negative regulator of the antiviral immune response,^34,35^ was also overall downregulated in rAAV9-treated males and females as well as in rAAV2-treated males (Figure 2A).

Conversely, *Ripk2*, involved in the activation of NF-κB and MAPK pathways with protective functions against viral infections,^36^ along with *Id2*, a negative regulator of antiviral immune responses,^37^ were upregulated following rAAV2 and rAAV9 treatment.

SnRNA-seq data revealed similar trends overall, with deregulation observed for both *Gadd45g* and *Irf2bp2* in rAAV2- and rAAV9-treated conditions (Figure S2A). Hepatocyte rather than immune cells were the primary source of expression of these genes (Figure S2B), which matched the absence of immune cell infiltrates observed in histopathology.

GSEA using the KEGG of ST data revealed significant downregulation of the complement and coagulation cascades in rAAV2- and rAAV9-treated females, along with antigen processing and presentation in periportal and intermediate zones (Figure S4B). Pathway activity predictions showed a significant downregulation of the hypoxia pathway in females, while EGFR and TGF-β pathways exhibited an overall upregulation trend in males and females (Figure 3F). These patterns were overall consistent in the snRNA-seq data (Figure S3B). These changes suggest a potential protective response to rAAV treatment, with partially sex-specific trends.

### ST delineates known and potential novel factors modulating rAAV entry

The zonation-associated distribution and transduction differences of rAAV9 and rAAV2 in the mouse liver may be linked to spatial variations in receptors, co-receptors, and other factors influencing rAAV transduction. Given the higher transgene capture efficiency of ST compared to snRNA-seq, we used the ST dataset to assess spatial factors, which potentially could modulate AAV entry and their association with EGFP transgene expression. It is currently unknown whether published AAV entry factors^38–42^ exhibit varying spatial expression patterns across liver zones or sex differences in mice, potentially modulating serotype- and sex-specific liver transduction patterns. Therefore, we first examined the zonal distribution and expression levels of known rAAV entry factors in control male and female mice.

Only *Fgfr1*, a known AAV2 co-receptor, showed higher expression in the portal zone of males compared to females (Figure S5), potentially contributing to the portal bias and sex difference in rAAV2 transgene expression. For other factors, either no zonation pattern was identified, or it did not correlate with the serotype-specific, periportal or pericentral zonation pattern or sex differences. snRNA-seq confirmed expression of all known entry factors in hepatocytes (Figure S5).

Next, we hypothesized that EGFP transgene expression would spatially correlate with the expression of known and putative novel AAV entry factors. Therefore, we analyzed transgene expression and its colocalization with membrane-associated genes as well as known AAV receptors (see Methods). Among the top spatially colocalized genes for rAAV9-treated males and females, we identified the gene *Rpsa*, encoding the laminin receptor, a known co-receptor for rAAV2 and rAAV9^41^ (Table S2). Additionally, the heparan sulfate proteoglycan (HSPG) *Sdc1* was found to colocalize with the transgene in rAAV2-treated males, consistent with the known role of HSPGs as AAV2 receptors.^43,44^ Further, *Itgb1* part of the α5β1 integrin, a coreceptor for rAAV2,^45^ was identified for the rAAV9-treated male animals. Finally, we also observed spatial co-expression of EGFP with *Dpp4,* which was recently described to bind to rAAV9.^40^

Top spatially colocalized genes for rAAV9-treated males and females (Figure 5A, B, Table S2) included the organic cation transporters *Slc22a1* and *Slco1b2*, as well as *Nrp1* and *Kdr*, the latter two have already been identified as a receptor for other viruses.^46,47^ Additionally, *Fcgrt*, encoding FcRn, a known target for therapeutic inhibition to reduce immune responses,^48^ was identified. In the ST data from control animals, none of the abovementioned genes showed consistent sex differences (Figure S6).

**Figure 5:**
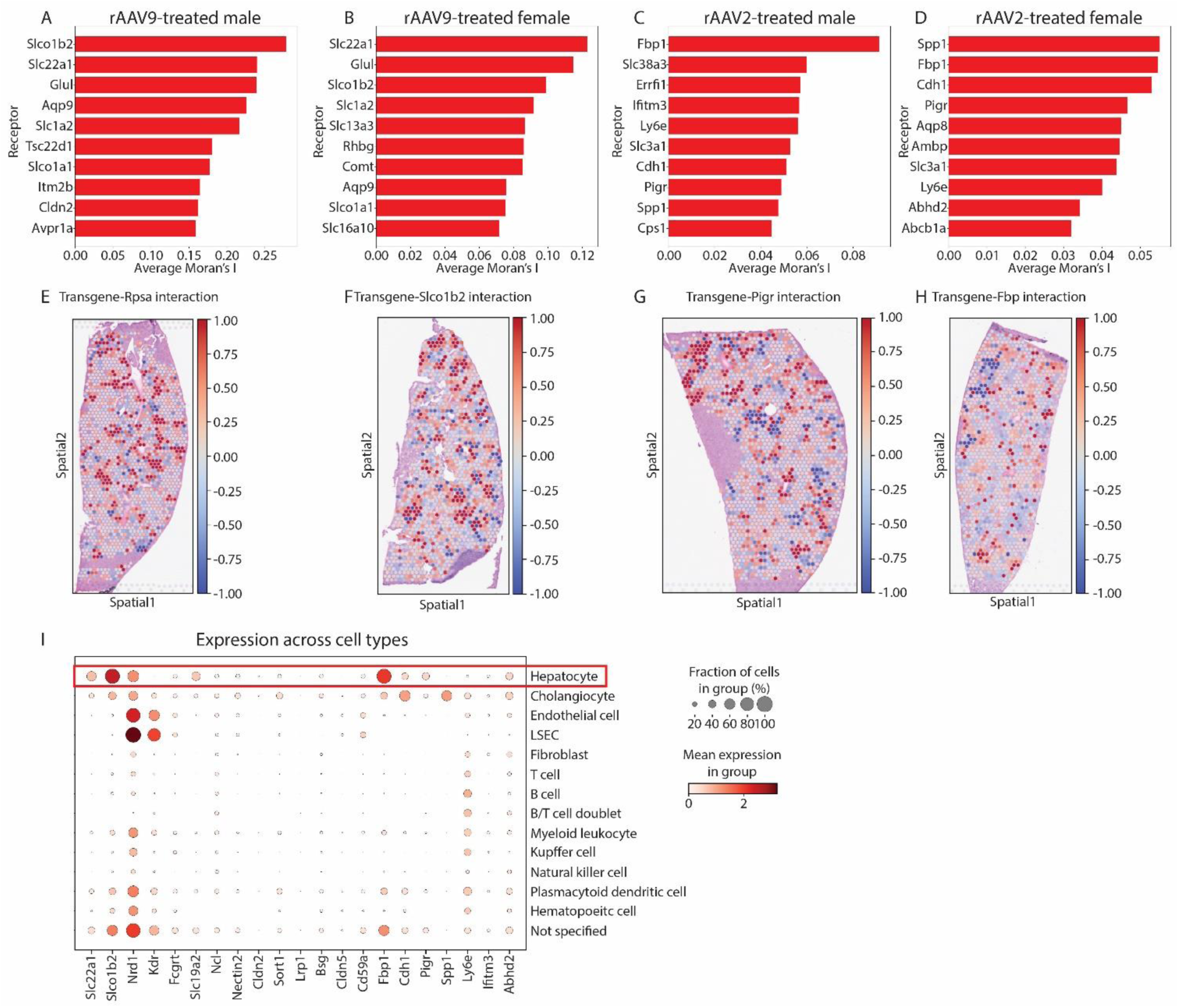
ST to investigate rAAV transgene and receptor colocalization. **(A-D)** Top 10 genes that show colocalization with the transgene in the spatial transcriptomics (ST) data for **(A)** rAAV9-treated males, **(B)** rAAV9-treated females. **(C)** rAAV2-treated males and **(D)** rAAV2-treated females. Genes are sorted by their average Moran’s I (a measure of spatial autocorrelation, see Methods). **(E-H)** Spatial mapping of the interaction between the transgene and **(E)** *Rpsa*, **(F)** *Slco1b2*, **(G)** *Pigr* and **(H)** *Fbp1* across a liver tissue section. **(I)** Cell type specific expression in the single-nucleus RNA sequencing (snRNA-seq) data combined for all animals. LSEC = liver sinusoidal endothelial cells.

Genes exclusively found in rAAV9-treated males (Figure 5A, Table S2) were *Slc19a2, Ncl*, and *Nectin2*, all of which have already been identified as entry receptors for other viruses or to be involved in promoting viral entry.^49,50,52,53^ *Cldn2* and claudins in general have been identified to regulate viral infection.^54^ *Sort1* and *Lrp1* were also identified, both involved in trafficking and endocytosis, respectively.^55–57^

Spatially colocalized genes exclusively observed in rAAV9-treated females (Figure 5B, Table S2) included *Bsg* linked to viral entry,^58,59^ and *Cldn5* and *Cd59a^60^* both implicated in regulating viral infections.

Among the top transgene colocalizing genes for rAAV2 in male and female mice (Figure 5C, D, Table S2) we identified *Fbp1, Cdh1, Pigr*, and *Spp1*, all of which are known to be involved in the entry of other viruses.^61–64^

Additionally, *Ly6e*, which promotes viral entry through mechanisms other than receptor-mediated entry,^65^ was also detected in rAAV2-treated livers of both sexes. The difference in expression between males and females in the ST data was also compared, and none of the transgene colocalizing genes showed significant consistent sex differences (Figure S6).

For rAAV2 males *Ifitm3* (Figure 5C, Table S2), known to negatively affect viral entry through mechanisms other than receptor-mediated entry,^66,67^ was identified.

In rAAV2 females *Abhd2* (Figure 5D, Table S2) was observed, which is known to promote viral propagation.^68^

Since ISH and IHC revealed transgene expression predominantly in hepatocytes, the expression of each of the above-mentioned cell membrane-expressed genes was examined per cell type (Figure 5I). Most of them were predominantly expressed in hepatocytes or also showed expression in other cell types besides hepatocytes. Exceptions were *Kdr* and *Cldn5*, which were mostly expressed in liver sinusoidal endothelial cells (LSECs) and *Spp1*, which seemed specific for cholangiocytes (Figure 5I).

Beyond receptors, we investigated factors recently emphasized by Rostami et al. for their roles in modulating AAV gene transfer and expression, analyzing their patterns in the context of sex differences and zonation. (Figure S7 and S8).^69^ None of the investigated genes showed differences that potentially could explain the observed sex and zonation differences.

In summary, our approach uncovered several established as well as novel factors that might influence rAAV uptake by hepatocytes and contribute to serotype- and sex-specific transduction patterns. This underscores the potential of ST to uncover intriguing targets for future research. These results shed light on both known and novel potential rAAV receptors, offering insights into the mechanisms of rAAV transduction and identifying potential targets for enhancing gene therapy efficacy.

## Discussion

This study highlights ST using the 10x Genomics Visium platform as a powerful tool to map AAV transduction patterns and reveals sex- and serotype-specific effects on gene expression in liver tissue. Visium ST offers unbiased, high-throughput transcriptomic profiling, enabling spatial resolution of transgene expression and tissue responses. When combined with snRNA-seq, it allows single-cell validation, overcoming the limitations of bulk RNA sequencing.

By exploiting ST and snRNA-seq, we systematically mapped rAAV2- and rAAV9-mediated transduction across hepatic zones and examined associated transcriptomic alterations.

Our findings uncover zonation-independent modulation of lipid metabolism, circadian rhythm, and immune responses, with several pathways displaying distinct sex-specific patterns. Moreover, our study identifies known and putative novel rAAV entry factors, providing insights into mechanisms underlying AAV tropism in hepatocytes. These findings highlight the potential of ST in gene therapy research, offering a comprehensive framework to study AAV vector biology *in situ*.

ST confirms that rAAV2 preferentially transduces periportal hepatocytes, while rAAV9 exhibits a bias toward pericentral regions, consistent with previous findings.^3^ Notably, we observed higher transgene expression in males, a phenomenon previously linked to androgen signaling.^5^ Beyond transduction efficacy, we identified profound transcriptomic changes influenced by sex and serotype, but largely independent of zonation. Some effects were consistent across both serotypes and sexes, indicating conserved mechanisms induced by rAAV transduction or transgene expression. Further experiments with empty capsid controls and promoterless transgenes are necessary, given EGFP’s reported cytotoxic effects.^70,71^

One of the most striking findings of our study is the serotype- and zonation-independent downregulation of lipid metabolism genes, including *Elovl3*, *Chka*, *Irs2*, and *Ppard*, across both sexes. These genes regulate lipid synthesis, transport, and beta-oxidation, suggesting that rAAV transduction broadly influences hepatic lipid homeostasis. Interestingly, *Srebf1*, *Tlcd4*, *Cpt2*, and *Acot1* showed sex-specific transcriptional changes, indicating that AAV vectors may differentially impact lipid metabolism in male and female mice. These findings align with recent studies linking viral infections to lipid metabolism dysregulation.^20,72–74^ Given that lipid metabolism is crucial for viral entry, replication, and immune evasion,^75–77^ our results raise the possibility that rAAV vectors may engage similar host pathways to optimize transgene expression. This insight is particularly relevant for AAV-based liver gene therapies, as pre-existing lipid metabolic conditions could influence vector efficacy.^78^

Moreover, to the best of our knowledge our study is the first to report that rAAV transduction alters hepatic circadian clock regulators. We observed zonation-independent upregulation of *Dbp, Tef*, *Nr1d1* and *Nr1d2* coupled with downregulation of *Arntl* (Bmal1) and *Nfil3*, suggesting that rAAV2 and rAAV9 vectors affect key circadian modulators. The circadian clock regulates hepatic metabolism, detoxification, and immune function, and its disruption has been linked to chronic liver diseases.^79^ The upregulation of *Dbp* is particularly interesting, as this gene contains a binding motif^80^ in the CMV enhancer, suggesting a potential feedback mechanism, where transgene expression influences host circadian regulation. Given that circadian disruption is known to affect viral replication and immune responses,^81–84^ these findings may have implications for gene therapy timing and efficacy.

Our transcriptomic analysis also uncovered immune regulatory genes that exhibit both serotype- and sex-specific expression patterns. In particular, *Gadd45g* —a regulator of replication stress and immune response^85,86^— was upregulated in males and downregulated in females following rAAV9 transduction. This sex difference may reflect variations in host immune surveillance mechanisms that influence AAV transduction efficacy. Additionally, we observed an upregulation of TGF-β pathway activity, indicating a potential pro-tolerogenic response to AAV transduction.^87^ The EGFR pathway was also upregulated, consistent with findings from a previous study utilizing an AAVrh.10 vector without a transgene,^88^ suggesting that the upregulation of EGFR may be attributed to viral transduction. Moreover, the downregulation of the hypoxia pathway in females may contribute to lower transgene expression in female mice due to hypoxia’s known role in episomal transcriptional regulation.^89,90^ In addition, the sex-independent suppression of *Rnf125* and *Irf2bp2*, genes modulating innate antiviral responses,^32–35^ indicate that AAV vectors may actively suppress immune activation. These findings have direct implications for gene therapy, particularly regarding sex-based differences in vector immunogenicity and persistence.

In addition to validating known AAV receptors (e.g. *Rpsa, Dpp4* for rAAV9, and *Sdc1*, for rAAV2), our ST analysis identified novel candidate receptors that may influence serotype-specific and sex-biased transduction patterns. These include solute carrier family proteins (e.g. *Slc22a1*, *Slco1b2*, *Slc19a2*), some of which have been implicated in viral uptake,^49^ as well as cell adhesion and trafficking proteins (e.g. *Cldn2*, *Nrp1*, *Kdr*), known to mediate virus-host interactions.^46,47,54^ Further functional validation (e.g. knockdown or overexpression studies) is required to confirm whether these factors directly modulate AAV transduction efficacy.

Despite its methodological strengths, our study has a few limitations. The sample size is limited (n=3 per condition), and future studies should expand sample numbers for better statistical power. Moreover, there are resolution constraints of Visium ST (55 μm per spot), which precludes single-cell resolution, but upcoming high-resolution platforms like Visium HD and Xenium may overcome this issue.^91^ In our study we also focused on one mouse strain, and future work should explore strain-specific effects on AAV transduction and host responses. Additionally, assessment of other time points post-transduction will allow for a more detailed investigation of the immune responses.^92,93^ Finally, our study design does not allow us to distinguish between transgene- or vector-induced effects, therefore, confirmation with empty vector controls is necessary. In regards to identifying potential novel rAAV entry factors, it is important to consider that mRNA and protein levels may not always correlate,^94,95^ which can hinder the identification of these receptors. Further studies using knockout models, receptor binding assays, and metabolic profiling may also deepen our understanding of AAV transduction mechanisms and their broader implications for gene therapy.

In conclusion, our study establishes ST as a powerful approach for investigating AAV transduction patterns and their molecular consequences in the liver. We reveal sex- and serotype-specific alterations in lipid metabolism, circadian regulation, and immune responses, and propose novel putative AAV entry factors. These findings provide critical insights into the host-vector interplay and highlight important considerations for optimizing AAV-based gene therapies. Future research should build on these findings to refine vector design and administration strategies, ultimately improving the safety and efficacy of gene therapy applications.

## Methods

### Mice

All studies were conducted with the approval of the cantonal veterinary authority of Basel-Stadt under license (BS-3082) followed in strict adherence to the Swiss federal regulations on animal protection. Mice were housed in groups of up to five in polycarbonate cages in a temperature (22 ± 2 °C) and humidity (55% ± 15%) controlled environment on a 12 h light cycle (06:00-18:00 h). The ventilation rate was more than 10 air exchanges per hour. Prior to study start mice are acclimatized for at least 7 days. 12 B6 Albino mice (B6N-Tyr^c-Brd^/Brd CrCrl) (6 male and 6 female) and 26 C57BL/6J mice (13 male and 13 female) of 6-8 weeks were supplied by either Charles River Laboratories (Saint-Germain-Nuelles, France) or Charles River Laboratories (Sulzfeld, Germany), respectively and randomly grouped (n=2-3) in IVC cages. Mice were housed with ad libitum access to food (Kliba Nafag 3439 pelleted) and water.

### rAAV vectors and administration

All the vectors (Table 1) were custom produced by SignaGen (Frederic, MD). For luminescence IVIS readouts, rAAV2 (Cat #SL100801) and rAAV9 (Cat #SL100801), carrying EGFP connected with the firefly luciferase (fLuc) via the Thosea asigna virus 2A peptide (T2A) (CMV-EGFP-T2A-fLuc), were delivered by the vendor at 1.10 x 1013 VG/ml and 3.68 x 1013 VG/ml concentrations respectively. For snRNAseq and histopathology, rAAV2 and rAAV9 carrying CMV-EGFP transgenes were delivered at 1.22 x 1013 VG/ml and 3.65 x 1013 VG/ml respectively. All batches were pre-tested for endotoxin and confirmed to contain less than 20% empty capsids. The EGFP transgene contains 70 CpG dinucleotides within its 720 nucleotides. For i.v. injection, formulation was done in sterile PBS (Cat #10010023, Gibco, Thermo Fisher Scientific, Carlsbad, CA) at 5 x 10^11^ VG per mouse. Mice were placed in the restrainer and 5 x 10^11^ VG were intravenously injected in 100 μl volume through the tail vein.

**Table 1:**
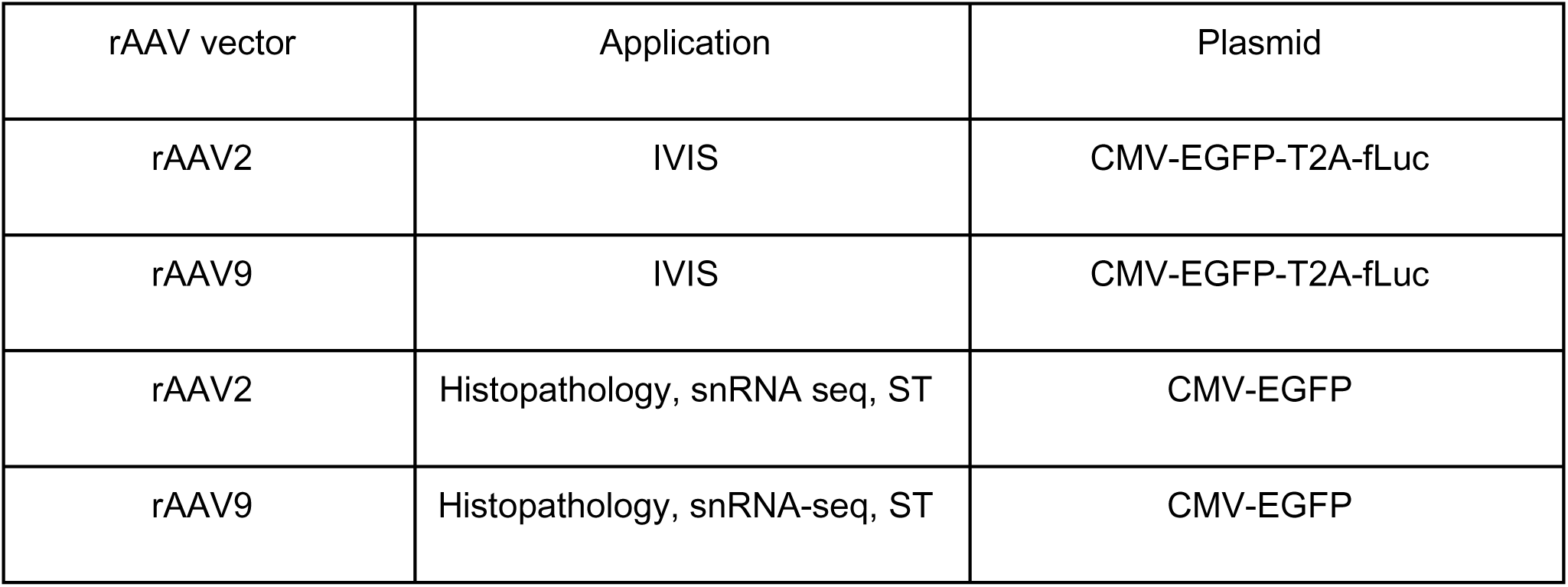
rAAV vectors used.

### IVIS

IVIS was performed on day 13, 22 and 27. The B6 albino mice were subdivided into groups of four and injected i.p. with IVISbrite™ D-Luciferin Ultra Bioluminescent Substrate in RediJect™ Solution (Cat #770505, Perkin Elmer, Waltham, MA) at 150 μg per gram of body weight. After 10 min, they were placed in an isoflurane induction chamber. The flow was maintained at 2.0 l/min at 4%. After 5 min the animals were placed inside the IVIS Spectrum system (Perkin Elmer) in a supine position, and the anesthesia was reduced to 2-3%. Subsequently, the images were acquired at FOV 22.8 cm (position D) using Auto Mode. The analysis was performed in Living Image software (Perkin Elmer). The elliptical ROIs were drawn in the liver area. Radiance values (p/s/cm/sr) calculated pixel-by-pixel were used to generate color-coded maps of luminescence, and corresponding Total Flux (p/s) values were reported in the graphs. Total Flux (p/s) values were calculated by summing radiance (p/s/cm²/sr) over the manually defined liver ROI using Living Image software.

### Tissue sampling

On day 28, animals were sacrificed and livers were collected. Parts of the left lateral lobe were either fixed in neutral buffered 10% formalin for 24 hours and processed for histology according to standard procedures or frozen in Optimal cutting temperature (OCT) using dry ice and stored at -80°C until further processing.

### Histopathology

#### Chromogenic ISH on fresh frozen and FFPE sections

Slides with 7 μm fresh frozen tissue sections were fixed in neutral buffered 10% formalin (Cat #9713.5000, VWR International, LLC., Randor, PA) for 15 min, followed by dehydration in increasing ethanol concentrations (50%, 70%, 2 x 100% 5 min each) at room temperature. The slides were air dried and subsequently transferred to the Ventana Discovery Ultra instrument (Ventana Medical Systems, Inc., Tucson, AZ, USA). mRNA expression of the targets was determined using the RNAscope VS Universal AP assay (Advanced Cell Diagnostics (ACD), Inc, Newark, CA) with the DISCOVERY mRNA RED, Amplification & Pretreatment PTO kit (Roche Diagnostics, Basel, CH, Cat #07074654001). ISH probes were all acquired from ACD and included EGFP (Cat #400289) as well as negative control DapB (Cat #312039) and the positive control Mm-PPIB (Cat #313919). The sections were counterstained with hematoxylin and bluing reagent for nuclei detection.

Formalin fixed, paraffin-embedded (FFPE) sections were cut at 3.5 μm and EGFP and EGFP-sense detection was performed using the RNAscope VS Universal HRP Detection reagents (Cat #323210, ACD), the Ventana Discovery mRNA Purple Detection Kit.(Cat #760-255, Ventana Medical Systems, Inc., Tucson, AZ, USA) and the Discovery mRNA Probe Amplification Kit (Cat #760-222, Ventana Medical Systems, Inc.). The sections were counterstained with hematoxylin and bluing reagent for nuclei detection.

#### IHC on FFPE sections

IHC was performed on FFPE liver tissue using the Ventana Discovery Ultra Instruments. Briefly slides were deparaffinized and antigen retrieval was performed using a citrate based puffer (pH 6.5) at 95°C for 32 min. EGFP was detected using a rabbit polyclonal GFP antibody (0.12 μg/ml, 37°C, 20 min, Cat #ab6556, Abcam Limited, Cambridge, UK) and an OmniMap anti-Rabbit HRP secondary antibody (Cat #760-4311, Ventana Medical Systems). Chromogenic detection was performed using the Discovery Purple Kit (Cat #760-229, Ventana Medical Systems). Sections were counterstained with hematoxylin and bluing reagent for nuclei detection.

#### Quantification of ISH and IHC on FFPE

Scanning was performed using a NanoZoomer S 360 C13220 series (Hamamatsu Photonics K.K., Hamamatsu City, Japan) at 40x magnification. The .ndpi image files were uploaded to HALO (Indica Labs, Albuquerque, NW). For evaluating percentage of positive cells in IHC and in ISH staining, first a pre-trained nuclear segmentation algorithm was used and fine-graded for our study. Subsequently, the ISH and IHC modules in HALO were adjusted to enhance their performance for their respective staining cases. After nuclear detection, the cytoplasm was set using nuclear extension. Quantification of the EGFP mRNA was performed using the cytoplasmic signal from the EGFP ISH probe. For EGFP protein the percentage of EGFP IHC positive was determined. In all the cases, data were segregated in (1) control, rAAV2, and rAAV9, and (2) in male, and female.

### Visium ST sample preparation

For RNA quality assessment, RNA from 3 animals per group was extracted using the Arcturus® PicoPure® RNA Isolation Kit (Cat #KIT0204, Applied Biosystems™, Waltham, MA). For cell lysis, two 10 μm sections of the sample were resuspended in 200 μl extraction buffer. Total RNA was extracted following the instructions of the manual. RNA integrity number (RIN) was assessed using the 2100 Bioanalyzer system (Agilent Technologies, Inc.) with an Agilent RNA 6000 Nano Kit (Cta # 5067-1511, Agilent Technologies, Inc.). Samples with RIN above 7.0 were used.

Tissue optimization was carried out according to the manufacturer’s instructions (Visium Spatial Tissue Optimization User Guide_RevC. 10x Genomics, Pleasanton, CA). Image acquisition was performed on the Hamamatsu NanoZoomer S 360 C13220 series (Hamamatsu Photonics K.K.) at 40x magnification and the coverslip was removed afterwards by immersing the slide in a 3x Saline-Sodium Citrate buffer (Cat #S66391L, Sigma-Aldrich, Merck KGaA, Darmstadt, Germany). The stained tissue sections were permeabilized using a time course to test for the optimal permeabilization time. After performing a fluorescent cDNA synthesis, the tissue was removed. Finally the fluorescent cDNA was imaged using a Zeiss Axio Scan.Z1 (Zeiss, Oberkochen, Germany) with a Plan Apochromat 20x/0.8 M objective, a ET-Gold FISH filter (ex 538-551 nm/em 556-560 nm) and 100 ms exposure time.

For the gene expression analysis, 10 μm thick sections of the samples were placed with a random distribution over chilled 10x Genomics Visium Gene Expression slides. From each group (control, rAAV2 and rAAV9) 3 males and females were included. The sections were similarly stained with H&E and subsequently imaged as described above. To release the mRNA, the sections were permeabilized for 12 min as defined by tissue optimization. For further processing, the cDNA was amplified according to the manufacturer’s protocol (CG000239_Visium Spatial Gene Expression User Guide_RevC). Double indexed libraries were prepared. The libraries were quality controlled using a 2100 Bioanalyzer system with Agilent High Sensitivity DNA Kit (Cat #5067-4626, Agilent Technologies, Inc.) and quantified with Qubit™ 1X dsDNA HS Assay Kit (Cat #Q33230, Invitrogen) on a Qubit 4 Fluorometer (Cat #Q33238, Invitrogen). The libraries were loaded onto the NovaSeq 6000 (Illumina, Inc., San Diego, CA) at a concentration of 150 pM. A NovaSeq SP v 1.5, S1 v 1.5 or S2 v 1.5 Reagent Kit (100 cycles) (Cat #20028319 and #20028401, Illumina, Inc) was used. For paired-end dual-indexed sequencing, the following read protocol was used: read 1: 28 cycles; i7 index read: 10 cycles; i5 index read: 10 cycles; and read 2: 90 cycles. All libraries were sequenced at a minimum of 50000 reads per covered spot.

Raw sequencing data were demultiplexed using the *mkfastq* function from Space Ranger (v. 1.2.0). Demultiplexed data were mapped to a custom reference with spaceranger count. The custom reference has been generated by adding the vector sequence to the mouse reference MM10 using spaceranger *mkref*. Spots under tissue folds, artifacts and tissue areas with insufficient morphological quality were manually removed using the 10x Genomics Loupe browser (v. 5.1.0).

### ST data analysis

#### Data preprocessing and quality control

SpaceRanger output was pre-processed and stored into anndata objects using the Scanpy^96^ and Squidpy^97^ packages. Upon examination of the overall spatial distribution of counts and detected genes, low quality spots were discarded following a similar approach to the one proposed by Ben Moshe and colleagues.^98^ In short, spots with total Unique Molecular Identifiers (UMI) count below two and above three Median Absolute Deviation (MAD) from the UMI count mean across all spots, computed individually for each sample, were filtered out. In the same line, spots with a fraction of mitochondrial genes greater than four MAD above the slide mean were also excluded from the analysis. Additionally, spots under tissue folds, artifacts, and those with insufficient morphological quality were removed. RNA counts per spot for each sample were normalized and log transformed. The *anndata* objects from the different samples were then concatenated, and Pearson’s Correlation coefficient was calculated between all the genes and the transgene.

#### Allocating Liver Zonation Regions

The pericentral and periportal regions of the mice liver were delineated by leveraging established markers.^2,8–10^ Specifically, gene expression of pericentral markers (*Glul, Cyp2e1, Oat, Slc1a2, Cyp1a2*) and periportal markers (*Sds, Cyp2f2, Hal, Hsd17b13, Alb, Arg1, Pck1*) was used to compute a per sample score using Scanpy’s *scanpy.tl.score_genes* function. Spots with scores in the top 80^th^ percentile of the per sample distribution were assigned to the corresponding pericentral or periportal category; the remaining spots were labeled as midzonal.

#### Deconvolution

We performed cell type deconvolution of the ST data using the snRNA-seq as the reference dataset. In our snRNA-seq reference, each nucleus was assigned a cell type label as described in the snRNA-seq analysis section. The nucleus annotated as *’B T cell doublet’* or *’mixed’* were discarded for this analysis. Cell type annotations were transferred from the snRNA-seq reference to each spot in the ST data by comparing spot-level expression profiles to those of the annotated single-nucleus reference. Deconvolution was carried out using two well-established methods, RCTD^11^ and cell2location^99^, to ensure that results were consistent across different approaches. Although both methods yielded similar estimates of the cell type composition, we present the findings from RCTD in the results section. The RCTD analysis was performed with default parameters and in ‘full mode’ to enable modeling of mixtures of multiple cell types per spot. The output of RCTD is a probability distribution of cell type mixtures for each spot.

For cell2location, we created the signature from our snRNA-seq data with the following parameters: *cell_count_cutoff = 25, cell_percentage_cutoff2 = 0.1*, and *nonz_mean_cutoff = 1.25*. This resulted in 8064 genes from 146663 nuclei on which the regression model was trained for 400 epochs using mice sex as a categorical covariate. Using this single-cell level signature, the cell2location model was then trained for 50000 epochs using the parameters *N_cells_per_location = 10 and detection_alpha = 100*. The final output of cell2location is a spot-by-cell-type matrix of estimated cell type abundances, which yielded similar results to the RCTD probabilities.

#### Pseudo-bulk and differential gene expression analysis

Raw counts from ST dataset were summed per sample for each liver region (pericentral, periportal, and midzone) to generate pseudo-bulk profiles using the decoupleR package^13^. The counts were then normalized to counts per million (CPM), log transformed, and scaled to a maximum of 10. Genes with fewer than 10 counts per sample or 40 reads across all samples were not included in the analysis.

Using PyDESeq2^100^ with Benjamini-Hochberg false discovery rate correction, differential gene expression analysis was performed between pseudobulk profiles for the different conditions (liver zone, AAV treatment, and sex). The apeglm shrinkage method^101^ was used to correct Log2 fold changes (log2FC), and differentially expressed genes were classified based on adjusted P values (p value<0.05) and absolute log2FC (>0.5) values.

#### Functional characterization

We estimated pathway activity per spot using PROGENy with a multivariate linear model as implemented in decoupleR (*decoupler.run_mlm*). The PROGENy model incorporates 14 pathways: Wnt, VEGF, Trail, TNFα, TGF-β, PI3K, p53, NFkB, MAPK, JAK/STAT, Hypoxia, Estrogen, Androgen, and EGFR. The model considers the expression of genes that are more responsive to perturbations on those pathways.^102^ In our analysis, we ran PROGENy using the top 500 most responsive genes for each pathway. Following the same procedure, the Wald statistic from the differential gene expression analysis on the pseudo-bulk profiles was used to estimate differences in pathway activities between conditions. A p value < 0.05 was considered significant for changes in pathway activity.

TFs activity per spot was determined using a univariate linear model (*decoupler.run_ulm*) on regulons derived from the CollecTRI resource.^103^ The Wald statistic from the differential gene expression analysis on the pseudo-bulk profiles was utilized to infer differences in TF activity between conditions using the same procedure. A p value < 0.05 was considered significant for changes in TF activity. The top 35 TFs with the most activity, regardless of condition, are displayed in the heatmap in the results section for comparison with the control.

The Gene Ontology Biological process (GO:BP)^14^ and Kyoto Encyclopedia of Genes and Genomes (KEGG)^15^ gene sets were used as reference to conduct GSEA on the differential gene expression analysis results on the pseudo-bulk profiles (*decoupler.get_gsea_df* with default parameters). Changes in a specific term (i.e., biological process or pathway) were considered significant if they show an adjusted p value < 0.05 after Benjamini-Hochberg correction for multiple hypothesis testing. The Scanpy *scanpy.tl.score_genes* function was employed to calculate a per spot score using the combined gene expression of *Nr1d1* and *Nr1d2* (selected for ‘RHYTHMIC_BEHAVIOR’), and *Eci2* and *Ech1* (selected for ‘FATTY_ACID_CATABOLIC_PROCESS’ and ‘FATTY_ACID_BETA_OXIDATION’). These genes were common leading-edge genes identified across tested conditions for these terms.

#### Inference of factors modulating rAAV entry

To investigate potential factors mediating rAAV entry into the cell, we employed LIANA+^104^ with the goal of identifying spatially co-expressed transgene-receptor pairs. To do this, we first defined the parameters of the spatial connectivity, specifically, by selecting a Gaussian kernel with a *bandwidth=150* and *cutoff=0.1*. We then computed the global Moran’s I score (a measure of spatial autocorrelation), applying 1000 permutations and considering 0.2 as the minimum expression proportion for receptors to be included in the analysis. As potential receptors, we included the corresponding genes of well-known receptors (from the literature) that mediate the entry of rAAV factors. These genes are *AU040320, Cd9, Fgfr1, Hspg2, Itgav, Itgb1, Itgb5, Met,* and *Rpsa*. Additionally, we extracted a list of receptors from CellCommuNet^105^ and, to obtain receptors relevant to the mouse liver, filtered by mus musculus, normal condition, single study type, and liver tissue. We also extracted potential receptors from UniProt,^106^ which are annotated as cell membrane genes (SL000039). It should be noted that the Moran’s I score is computed per sample; therefore, we subsequently averaged the scores for the samples belonging to the same condition. We considered relevant genes to be those receptors with an average per condition Moran’s score above 0.025 (with the transgene) and a permutation-based p value below 0.01. Finally, and for visualization purposes, we selected the top 10 receptors exhibiting the largest global Moran’s score with the transgene.

### Nuclei isolation

Nuclei were isolated from fresh frozen liver samples. Samples from 3 control males and females and 4 rAAV2- or rAAV9-treated males and females were processed, respectively. One rAAV9-treated female was processed in two replicates. Tissue samples of 17-35 mg were dissociated using a Singulator 100 (S2 Genomics, Inc., Livermore, CA) Nuclei suspensions were centrifuged at 500 g for 5 min at 4°C and resuspended in Nuclei Storage Reagent (Cat #100-058-874, S2 Genomics, Inc.) supplemented with 0.2 U/ul RNaseOUT™ Recombinant Ribonuclease Inhibitor (Cat #10777019, Thermo Fisher Scientific). Nuclei were stained with NucBlue™ Fixed Cell ReadyProbes™ Reagent (Cat #R37606, Thermo Fisher Scientific) containing 4’,6-diamidine-2’-phenylindole dihydrochloride (DAPI) at one drop per 500 μl. After 30 min incubation on ice cells were strained through 35 μm cell strainers prior to fluorescence-activated cell sorting (FACS) using a FACSAria™ Fusion Flow Cytometer (BD Biosciences, San Jose, CA, USA). Cell strainers were pre-wet with 1% BSA in PBS. Nuclei were identified by size and signal intensity. Nuclei were sorted into a pre-coated tube containing 1% bovine serum albumin (BSA, Cat #A9576-50mL, Sigma) in PBS. Subsequently nuclei were centrifuged at 500 g for 5 min at 4°C and resuspended in PBS containing 0.04% BSA using pre-coated filter tips. Nuclei were manually counted using a hemocytometer and used for single-nucleus RNA sequencing.

### Single-nucleus RNA sequencing

For the construction of snRNA-seq libraries, Chromium Next GEM Single Cell 3’ Kit v3.1 (10x Genomics, Pleasanton, CA) was used according to the manufacturer’s instructions. 8,000 sorted nuclei were loaded onto a 10x Genomics Chromium Next GEM Chip G and processed immediately in a 10x Genomics Chromium controller. Amplification of cDNA was performed during 12 PCR cycles. Libraries were quantified with a Qubit™ 1X dsDNA HS Assay Kit (Cat #Q33230, Invitrogen, Waltham, CA) on a Qubit 4 Fluorometer (Cat #Q33238, Invitrogen). Fragment size was assessed with a TapeStation High Sensitivity D5000 Screen Tape (Agilent Technologies, Inc., Santa Clara, CA). Libraries were pooled and loaded at a concentration of 200 pM and sequenced using a NovaSeq 6000 (Illumina, Inc., San Diego, CA) and S4 flow cell (Cat #20028313, 200 cycle kit) with a read one length of 28 cycles, and a read two length of 90 cycles. All libraries were sequenced at a minimum of 25000 reads per covered cell. Raw sequencing data were demultiplexed using the *mkfastq* function from CellRanger (v. 6.0.2 10x Genomics). Using the CellRanger *count* Demultiplexed data were mapped to a custom reference including the vector sequence generated by *mkref* function.

### snRNA-seq data analysis

#### Data preprocessing and quality control

CellBender, a variational inference framework used to estimate and remove ambient RNA contamination from droplet-based single-cell (or snRNA-seq) data, was employed to mitigate technical artifacts and ambient RNA signals prior to downstream analysis.^107^ CellBender was employed with default parameters for *fpr* (0.01) and *epochs* (150) for all the samples, while the expected number of cells and the *learning-rate* were adjusted for each sample to prevent over-correction.

Post–ambient RNA removal, nuclei were subjected to a stringent filtering process based on unique molecular identifiers (UMIs), number of detected genes, and percentage of mitochondrial transcripts. Specifically, we excluded nuclei with < 1000 counts or > 40000 counts, < 400 genes or > 8000 genes and > 1% mitochondrial reads.

#### Downstream analysis and cell type annotation

The Besca toolkit, an automated single-cell analysis pipeline implemented in Python and built upon Scanpy, was employed for clustering and subsequent interpretative analyses.^96,108^ Besca streamlines typical sc/snRNA-seq analysis steps, including normalization, feature selection, dimensionality reduction, clustering, and cell type annotation. Briefly, RNA counts were normalized per 10000 (cp10k), the top highly variable genes were selected for each sample, and a principal component analysis (PCA) was performed. The first 50 principal components (PCs) were used for nearest neighbor (*n_neighbors = 10*) calculations and Leiden clustering (*resolution = 1.5*). To visualize the data, uniform manifold approximation and projection (UMAP) embeddings were computed from the PCA space.

Cell annotation was performed using Besca’s cell annotation workflow and the *sig-annot* module, with the nomenclature following cell ontology (CL) conventions wherever possible. Cell type annotation was performed using Besca markers, along with additional markers detailed in Table S3. The UMAP in Figure S1G shows a coarse-grained version of this annotation (Table S4). The detailed cell type annotations from Table S3 were used for deconvolution.

#### Pseudo-bulk and differential gene expression analysis

Raw counts from the snRNA-seq dataset were summed independently for each sample, for both periportal and pericentral annotated hepatocytes, to generate pseudo-bulk profiles using the decoupleR package.^13^ The counts were then normalized to counts per million (CPM), log transformed, and scaled to a maximum of 10. Genes with fewer than 10 counts per sample or 15 reads across all samples were excluded from the analysis.

Differential gene expression analysis was performed between pseudobulk profiles following the same procedure described above for the ST data.

#### Functional characterization

As described in the corresponding ST section, the Wald statistic from the differential gene expression analysis on the pseudo-bulk profiles of pericentral and periportal hepatocytes was used to estimate differences in pathway and transcription factor activities between conditions. A p value < 0.05 was considered significant for changes in pathway or transcription factor activity.

GSEA was performed on the differential gene expression analysis results derived from the pseudo- bulk profiles of pericentral and periportal hepatocytes. This analysis followed the same methodology as described for the ST dataset. Significant changes within a specific term (e.g., biological process or pathway) were determined using a Benjamini-Hochberg adjusted p value threshold of less than 0.05 to account for multiple hypothesis testing.

### Statistical analysis

Statistical methods for each analysis are detailed in their respective sections. Pearson correlation coefficients were computed using the *numpy.corrcoef* function from the NumPy Python package. We compared average gene expression levels between male and female groups across zonation regions within each experimental condition by performing the Welch’s t-test (*scipy.stats.ttest_ind*).

The figures, including heatmaps, spatial maps, scatterplots plots, bar plots and dot plots were created using Matpotlib,^109^ Scanpy and Seaborn.^110^ Dotplots to show quantification of ISH, IHC and snRNA-seq were generated using GraphPad Prism v10.1.2 (GraphPad Software, Boston, Massachusetts USA).

## Data and code availability

The output of Spaceranger, including processed count data matrices and histological images, for the ST data generated in this study is available at https://zenodo.org/records/14917686.

In addition, this repository also contains the cellranger outputs as well as the CellBender output. The BAM files from the snRNA-seq data are available upon request. The scripts containing all the code used to generate the results presented in this study are available at https://github.com/alberto-valdeolivas/ST_SN_AAV_MiceLiver.

## Acknowledgements

This work was supported by the ARDAT consortium. We thank Virginie Ott for the support with the EGFP ISH and IHC.

## Author contributions

Conceptualization: A.V., P.C.S. and K.H.; Investigation: B.A., F.K. F.B., L.S.; Data curation and formal analysis: B.A., M.H.O., F.B., P.C.S., A.V. and K.H.; Supervision: N.K., M.P., M.B.O., F.S., A.N., D.M., H.H., R.X., M.S., B.J., S.R.; Writing - original draft: B.A. and K.H.; Writing – review & editing: all authors.

## Declaration of interests

N.K., M.B.O., H.H., R.X., M.S., B.J., A.V., P.C.S. and K.H. are shareholders of F. Hoffmann-La Roche Ltd. A.N. is a shareholder of Spark Therapeutics Inc a subsidiary of Roche. All other authors declare no competing interests.

## Notes

https://zenodo.org/records/14917686

https://github.com/alberto-valdeolivas/ST_SN_AAV_MiceLiver

### Competing Interest Statement

B.A., F.K., M.P., M.H.O., and F.S. are employees of F. Hoffmann-La Roche Ltd.. N.K., M.B.O., H.H., R.X., M.S., B.J., A.V., P.C.S. and K.H. are employees and shareholders of F. Hoffmann-La Roche Ltd.. A.N. is an employee and shareholder of Spark Therapeutics Inc a subsidiary of Roche. D.M. is an employee of Spark Therapeutics Inc. All other authors declare no competing interests.

### Summary of Updates

The abstract and the graphical abstract were revised. Headings of Figure 2-4 were rephrased for better clarity.

