## Supplementary Figures S1-S8 and Supplementary Tables S1, S3, S4 for "Spatial Transcriptomics and Single-Nucleus RNA Sequencing Reveal rAAV2- and rAAV9-Specific Transduction Signatures in the Mouse Liver"

* Corresponding author

### Supplemental Figures

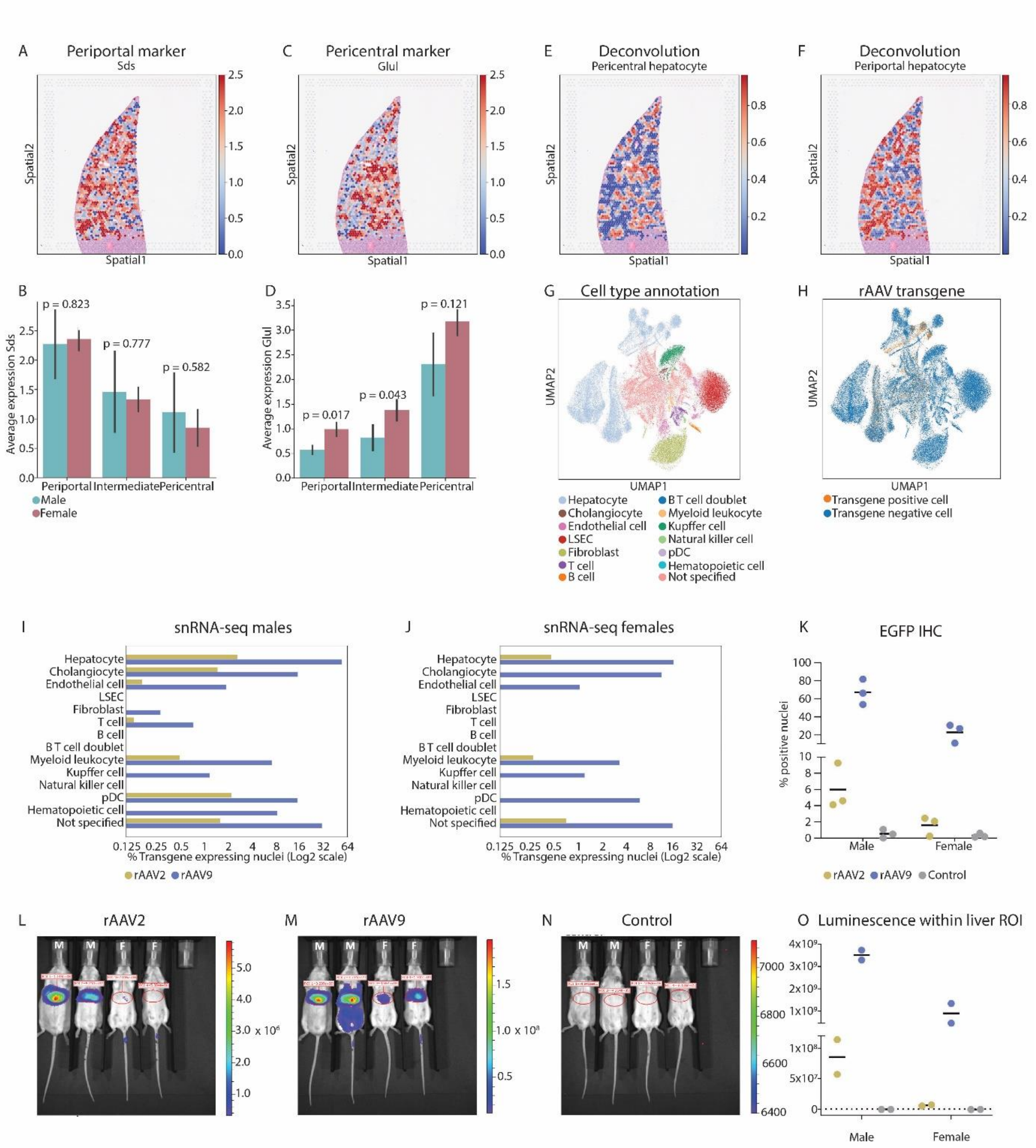

**Figure S1: Transgene distribution in rAAV2- and rAAV9-treated mice. (A)** Spatial mapping of the periportal marker *Sds* across the liver of a rAAV9-treated male mouse. **(B)** Average expression of *Sds* in the liver zones of the control animals. Black bars represent the standard deviation per condition. Statistical analysis to compare males and females was performed using Welch's t-test (Methods), P values are indicated. **(C)** Spatial mapping of the pericentral marker *Glul* across the liver of a rAAV9-treated male mouse. **(D)** Average expression of *Glul* in the liver zones of the control animals. Black bars represent the standard deviation per condition. Statistical analysis to compare males and females was performed using Welch's t-test (Methods), P values are indicated. **(E)** Spatial mapping of the predicted percentage of periportal hepatocytes per spot, using deconvolution. **(F)** Spatial mapping of the predicted percentage of pericentral hepatocytes per spot, using deconvolution. **(G)** UMAP embedding of cell clusters from single-nucleus RNA sequencing (snRNA-seq) with annotations. LSEC = liver sinusoidal endothelial cell, pDC = plasmacytoid dendritic cell. **(H)** snRNA-seq UMAP embeddings of the gene expression measurements colored for the transgene expressing nuclei. **(I)** Percentage of transgene-expressing nuclei per cell type in male mice treated with rAAV2 and rAAV9 in the snRNA-seq data. Percentages are normalized to each respective treatment condition and plotted on a Log2 scale. **(J)** Percentage of transgene-expressing nuclei per cell type in female mice treated with rAAV2 and rAAV9 in the snRNA-seq data. Percentages are normalized to each respective treatment condition and plotted on a Log2 scale. **(K)** Liver tissue immunohistochemistry (IHC) for EGFP protein expression. Graph shows quantification of IHC as percent of positive cells for male and female rAAV2-, rAAV9-treated and control animals. Each dot represents one individual with the mean represented by horizontal lines. **(L-O)** Bioluminescence images from an In Vivo Imaging System (IVIS. Methods) showing the transgene expression in male (M) and female (F) B6 Albino mice, 27 days post intravenous injection of **(L)** rAAV2 and **(M)** rAAV9 with a CMV-EGFP-T2A-fLuc transgene or **(N)** PBS for control animals. Color bars denote radiance (p/sec/cm$^2$/sr). **(O)** Quantification of the IVIS fluorescent signal in the liver. Total Flux (p/s) values were calculated by summing radiance (p/s/cm$^2$/sr) over the manually defined liver region of interest (ROI) (K, L, M) using Living Image software. Each dot represents one individual with the mean represented by horizontal lines. The dashed gray line corresponds to the background signal level in control animals.

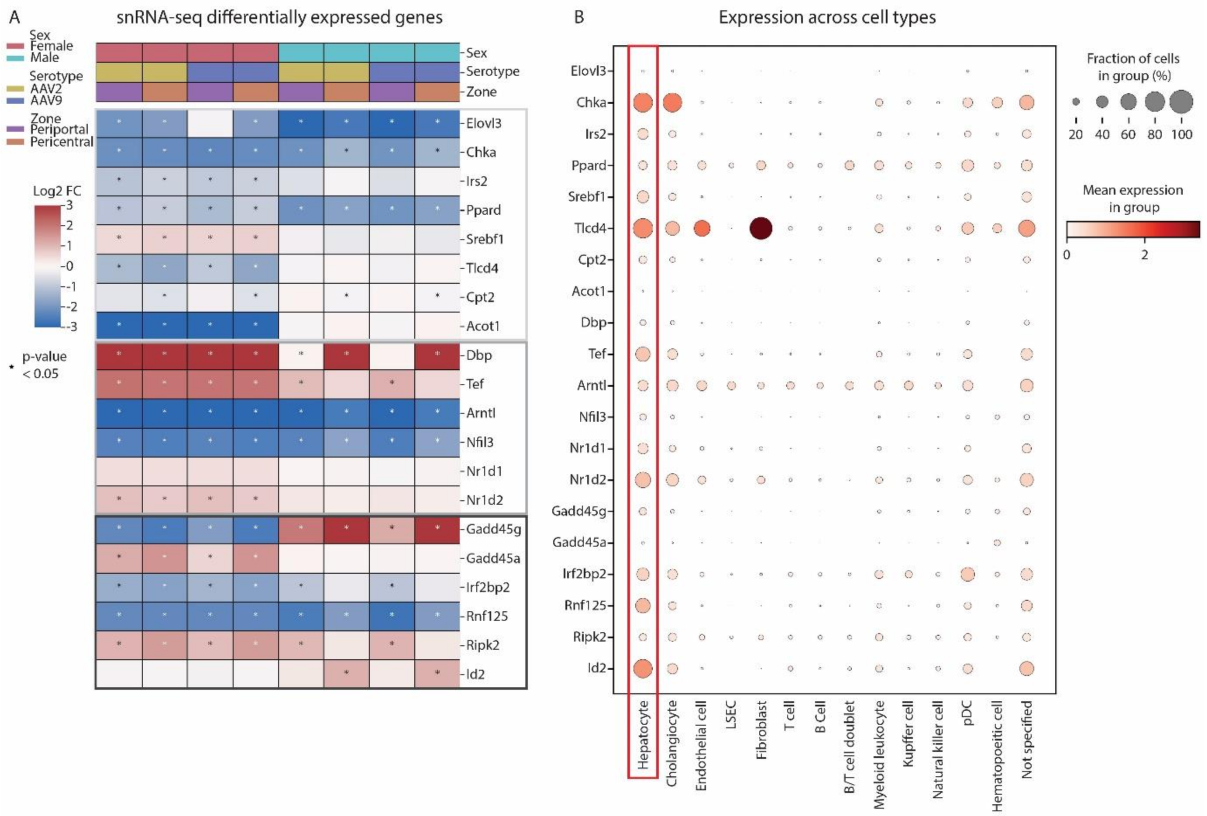

**Figure S2: SnRNA-seq confirms changes in DEGs and enables cell-type specific assessment of expression. (A)** Differential gene expression in treated versus control animals, categorized by liver zone and sex in the single-nucleus RNA sequencing (snRNA-seq data). Displayed is a selection of genes related to lipid metabolism (light grey square), circadian rhythm (medium gray square), and immune modulation (dark grey square). P values were calculated using the Wald statistical test (Methods). **(B)** Average expression of discussed genes across the cell types in all the samples in the snRNA-seq data. LSEC =liver sinusoidal endothelial cell, pDC = plasmacytoid dendritic cell.

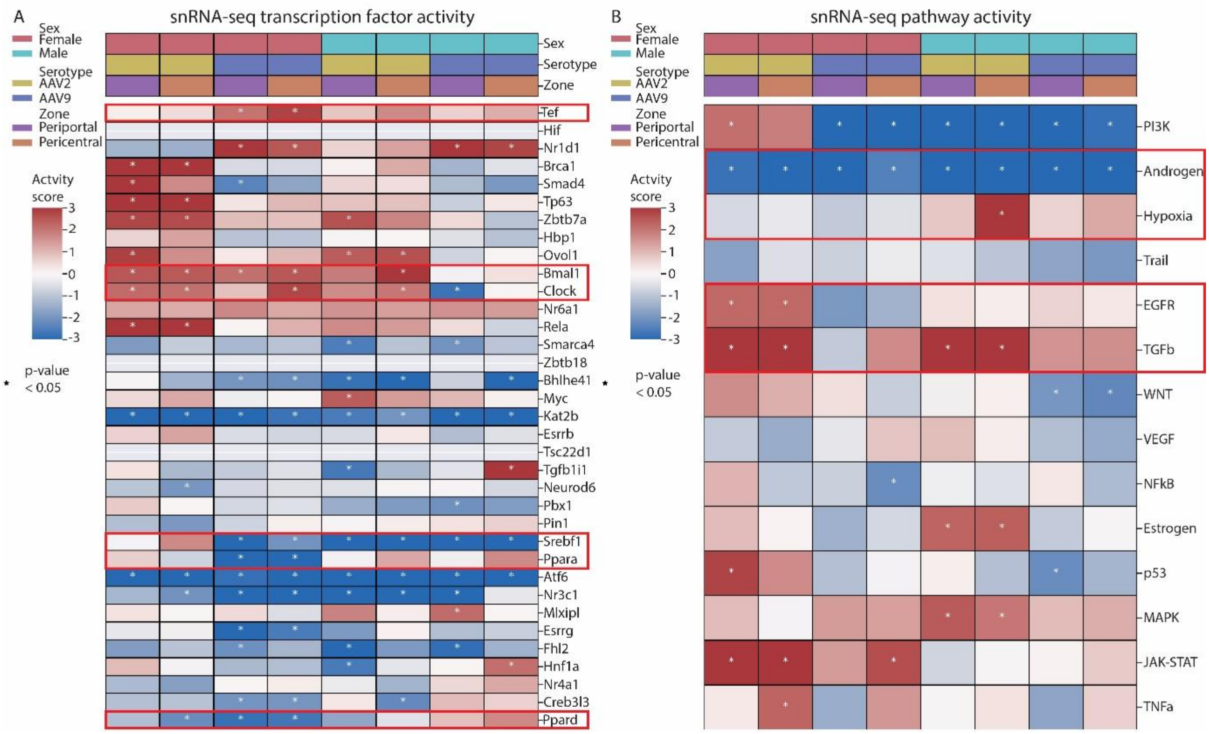

**Figure S3: snRNA-seq confirms changes in transcription factor and pathway activity. (A)** Predicted transcription factor activity in the periportal and pericentral hepatocytes in the single-nucleus RNA sequencing (snRNA-seq) data. P values were calculated using the Wald statistical test (Methods). Transcription factor activity for Hif, Zbtb18 and Tsc22d1 could not be predicted in the snRNA-seq data. **(B)** Differential pathway activity computed on pseudo-bulk RNA-seq generated from the periportal and pericentral annotated hepatocytes in the snRNA-seq data. P values were calculated using the Wald statistical test (Methods). The transcription factors and pathways discussed in the text are highlighted with a red square.

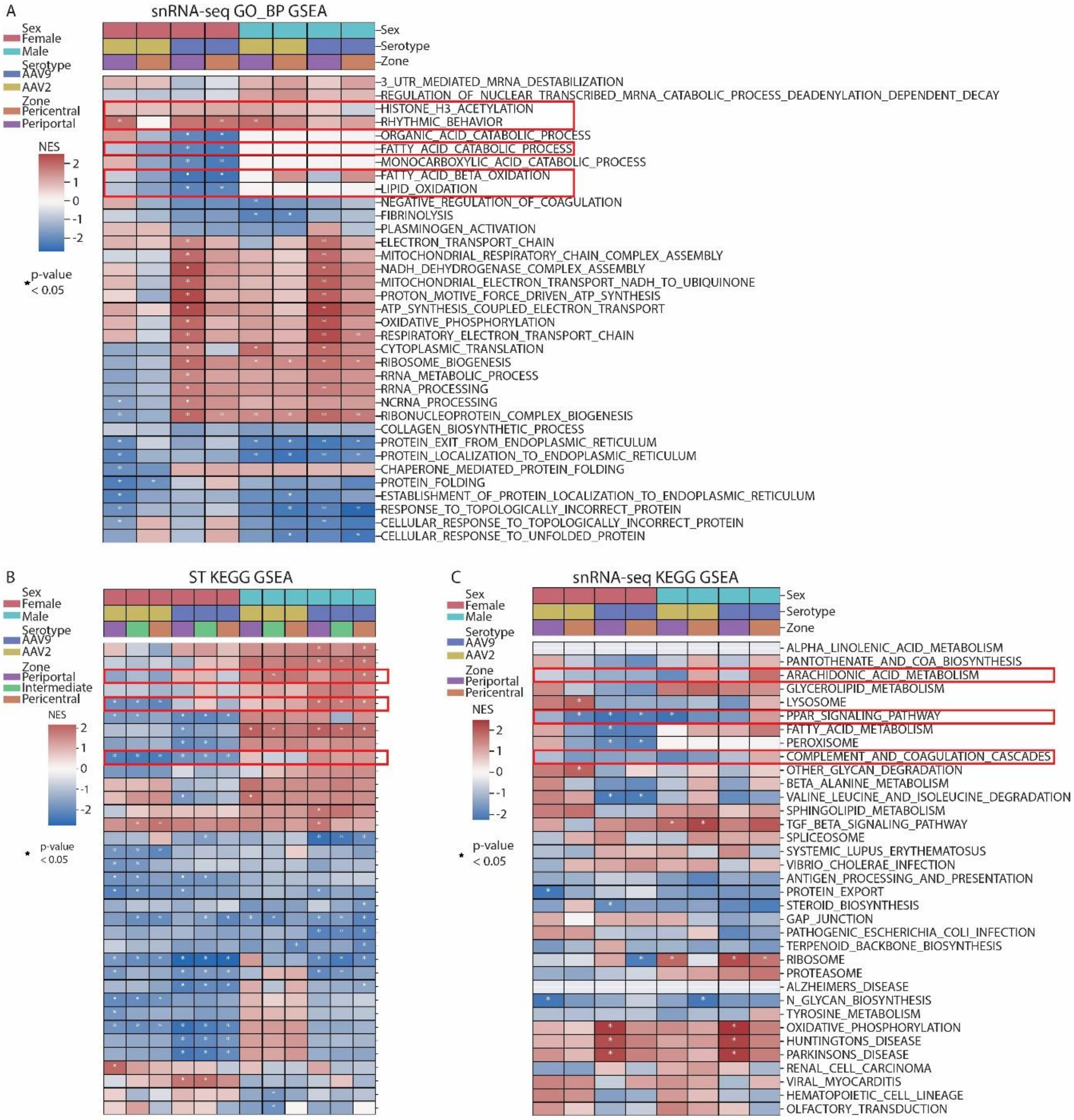

**Figure S4: GSEA in ST and snRNA-seq data. (A)** Gene set enrichment analysis (GSEA) using the Gene ontology biological processes (GO_BP) database in the single-nucleus RNA sequencing (snRNA-seq) data. **(B)** GSEA using the Kyoto Encyclopedia of Genes and Genomes (KEGG) database in the spatial transcriptomics (ST) data. **(C)** GSEA using the KEGG database in the snRNA-seq data. The analysis did not produce results for ‘ALPHA_LINOLENIC_ACID_METABOLISM‘ and ‘ALZHEIMERS_DISEASE‘ on the snRNA-seq data because the thresholding requirements were not met. Gene sets discussed in the text are highlighted with a red square. P values were calculated using the Wald statistical test (Methods).

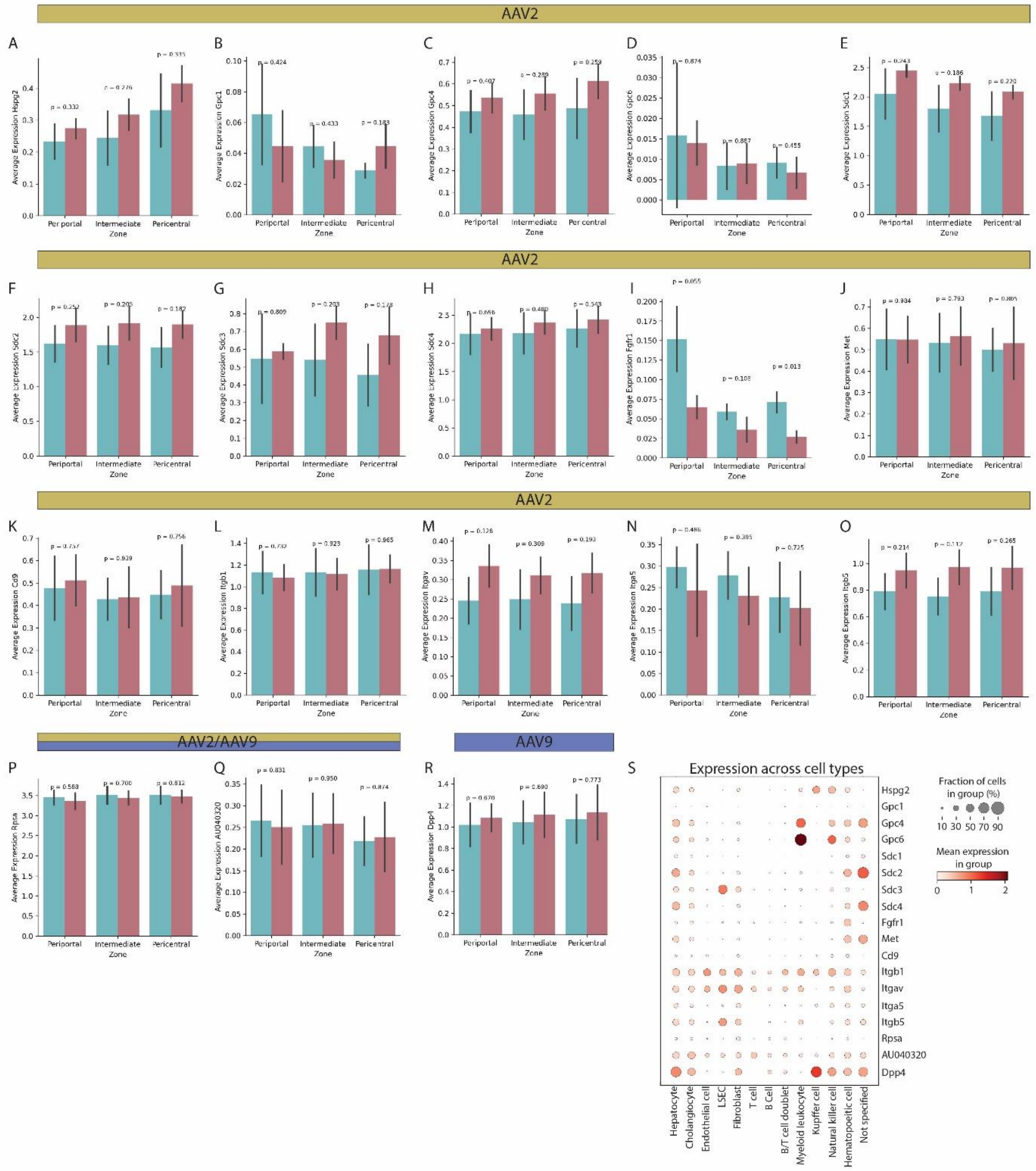

**Figure S5: Zonal distribution of known rAAV2 and rAAV9 receptors in male and female control mice. (A)** *Hspg2*, **(B)** *Gpc1*, **(C)** *Gpc4*, **(D)** *Gp6*, **(E)** *Sdc1*, **(F)** *Sdc2*, **(G)** *Sdc3*, **(H)** *Sdc4*, **(I)** *Fgfr1,* **(J)** *Met*, **(K)** *Cd9*, **(L)** *Itgb1*, **(M)** *Itgav*, **(N)** *Itga5*, **(O)** *Itgb5*, **(P)** *Rpsa*, **(S)** *AU040320,* **(R)** *Dpp4*. Data is taken from the spatial transcriptomics data. Black bars represent the standard deviation per condition. Statistical analysis to compare males and females was performed using Welch's t-test (Methods), P values are indicated. **(S)** Expression of the receptors across cell types of the control animals in the single-nucleus RNA sequencing (snRNA-seq) data. LSEC = liver sinusoidal endothelial cells. Plasmacytoid dendritic cells are not displayed as they were not identified in the control animals.

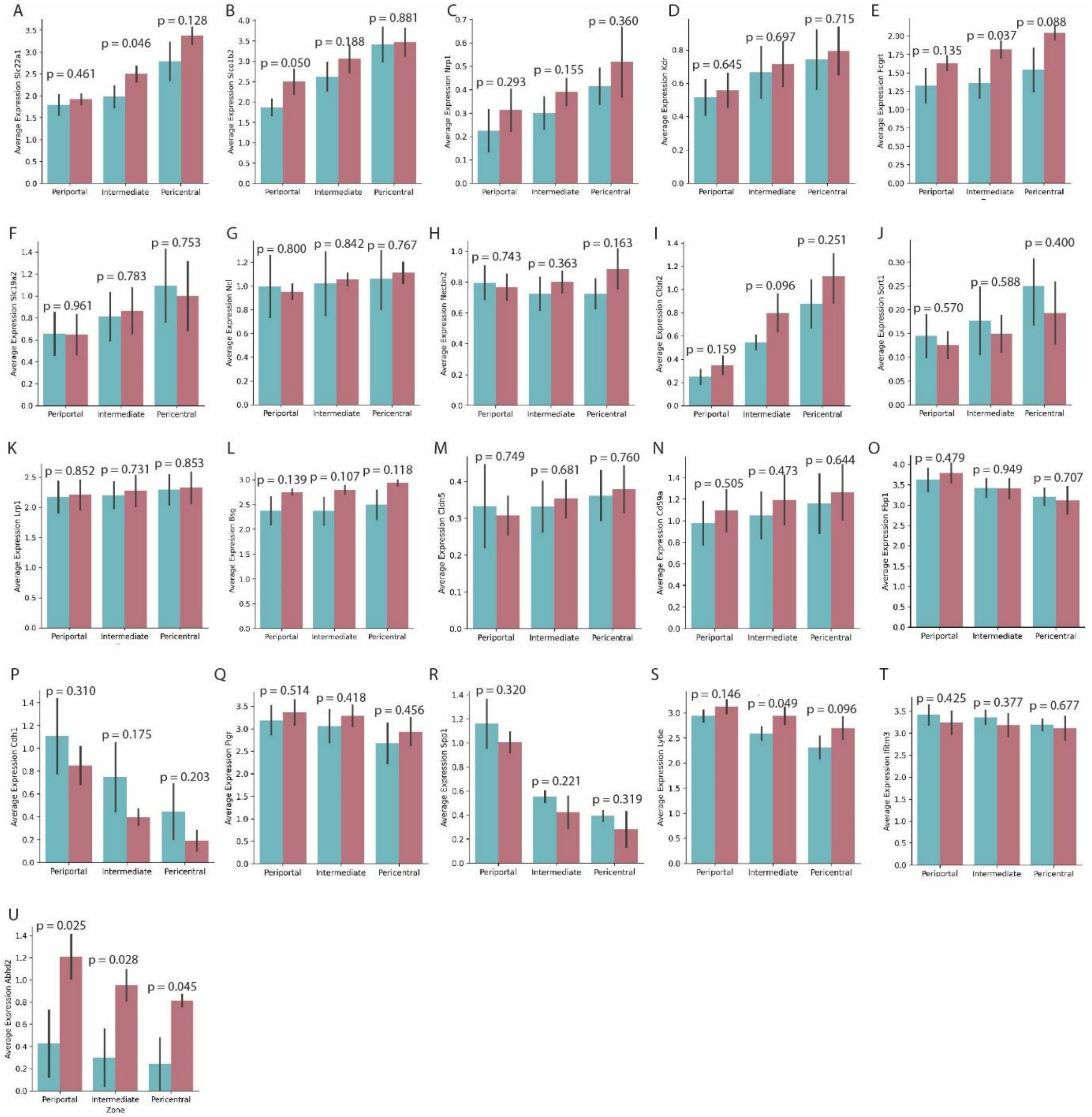

**Figure S6: Zonal distribution of potential novel AAV-entry factors in males and females in the control animals in the ST data. (A)** *Slc22a1*, **(B)** *Slco1b2*, **(C)** *Nrp1*, **(D)** *Kdr*, **(E)** *Fcgrt*, **(F)** *Slc19a2*, **(G)** *Ncl*, **(H)** *Nectin2*, **(I)** *Cldn2*, **(J)** *Sort1*, **(K)** *Lrp1*, **(L)** *Bsg*, **(M)** *Cldn5*, **(N)** *Cd59a*, **(O)** *Fbp1*, **(P)** *Cdh1*, **(Q)** *Pigr*, **(R)** *Spp1*, **(S)** *Ly6e*, **(T)** *Ifitm3*, **(U)** *Abhd2*. Data is taken from the spatial transcriptomics (ST) data. Black bars represent the standard deviation per condition. Statistical analysis to compare males and females was performed using Welch's t-test (Methods), P values are indicated.

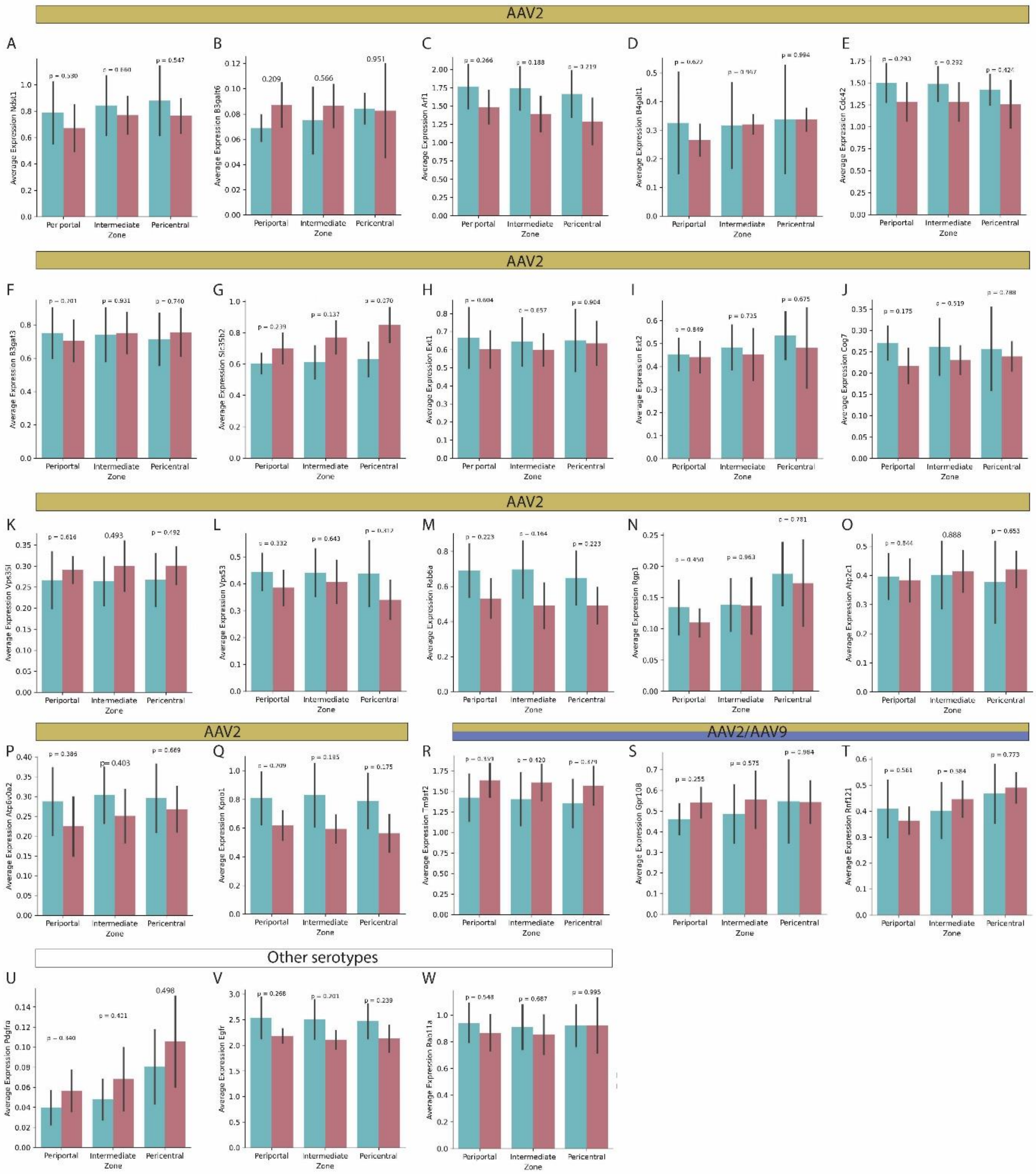

**Figure S7: Zonal distribution of factors known to positively affect AAV transgene expression in male and female control mice. (A)** *Ndst1*, **(B)** *B3galt6*, **(C)** *Arf1*, **(D)** *B4galt1*, **(E)** *Cdc42*, **(F)** *B3gat3*, **(G)** *Slc35b2*, **(H)** *Ext1*, **(I)** *Ext2*, **(J)** *Cog7*, **(K)** *Vps35l*, **(L)** *Vps35*, **(M)** *Rab6a*, **(N)** *Rgp1*, **(O)** *Atp2c1*, **(P)** *Atp6v0a2*, **(Q)** *Kpnb1,* **(R)** *Tm9sf2*, **(S)** *Gpr108*, **(T)** *Rnf121*, **(U)** *Pdgfra*, **(V)** *Egfr*, **(W)** *Rab11a*. Data is taken from the spatial transcriptomics data. Displayed are the control animals. Black bars represent the standard deviation per condition. Statistical analysis to compare males and females was performed using Welch's t-test (Methods), P values are indicated.

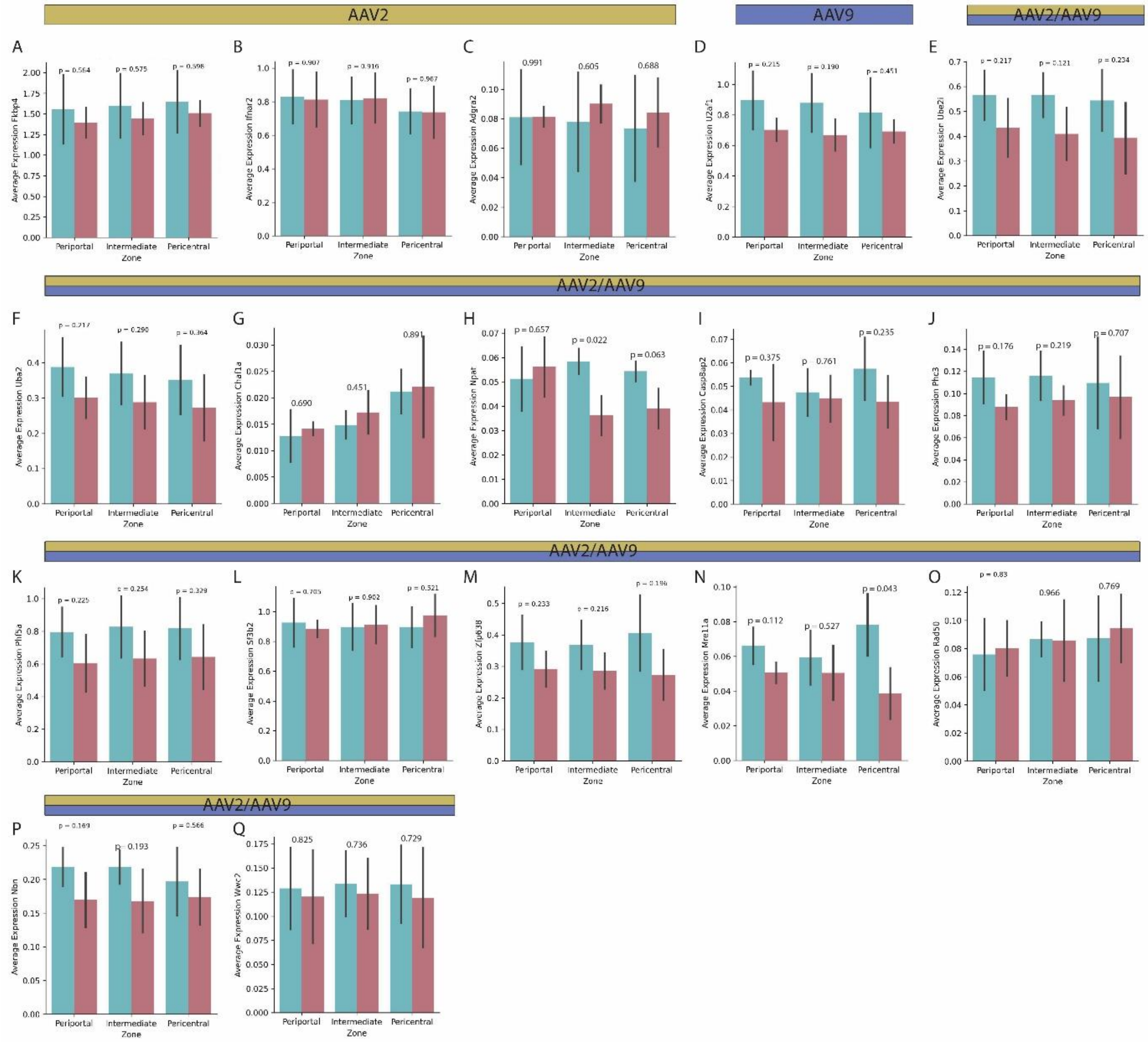

**Figure S8: Zonal distribution of factors known to negatively affect AAV transgene expression in male and female control mice. (A)** *Fkbp4*, **(B)** *Ifnar2*, **(C)** *Adgra2*, **(D)** *U2af1*, **(E)** *Ube2i*, **(F)** *Uba2*, **(G)** *Chaf1a*, **(H)** *Npat*, **(I)** *Casp8ap2*, **(J)** *Phc3*, **(K)** *Phf5a*, **(L)** *Sf3b2*, **(M)** *Zfp638*, **(N)** *Mre11a*, **(O)** *Rad50*, **(P)** *Nbn*, **(Q)** *Wwc2*. Data is taken from the spatial transcriptomics data. Displayed are the control animals. Black bars represent the standard deviation per condition. Statistical analysis to compare males and females was performed using Welch's t-test (Methods), P values are indicated.

### Supplemental Tables

**Table S1: Cell type distribution among the transgene positive cells.** Cell type proportions were determined as percentages among the transgene positive cells, categorized per serotype and sex in the single-nucleus RNA sequencing data.

| Cell type | rAAV2-CM-GFP_FeMale | rAAV2-CMV-GFP_Male | rAAV9-CMV-GFP_Female | rAAV9-CMV-GFP_Male |
|---|---|---|---|---|
| Kupffer cell | 0.00 | 0.38 | 0.34 | 0.28 |
| LSEC | 0.00 | 0.58 | 0.00 | 0.00 |
| T cell | 0.00 | 0.19 | 0.00 | 0.07 |
| Cholangiocyte | 0.00 | 0.77 | 0.41 | 0.62 |
| ndothelial cell | 0.00 | 0.96 | 0.11 | 0.23 |
| Fibroblast | 0.00 | 0.38 | 0.05 | 0.20 |
| matopoietic cell | 0.00 | 0.00 | 0.00 | 0.02 |
| Hepatocyte | 58.33 | 51.06 | 67.99 | 60.07 |
| eloid leukocyte | 0.76 | 0.58 | 0.30 | 0.70 |
| Not specified | 40.91 | 43.38 | 30.64 | 37.20 |
| cytoid dendritic cell | 0.00 | 1.73 | 0.16 | 0.60 |

**Table S2: Genes that show colocalization with the transgene.** Genes are sorted by their average Moran's I (see Methods). Genes discussed in the text are highlighted in red. (Excel file)

**Table S3: Cell type markers and signatures used for annotation of snRNA-seq data.**

| Cell type | Marker | | | | | | | | | | |
|---|---|---|---|---|---|---|---|---|---|---|---|
| Periportal hepatocyte | Pck1 | Ttr | Fabp1 | Tat | Sds | Hal | Cyp2f2 | Hsd17b13 | Asl | Ass1 | |
| Hepatocyte | Pck1 | Ttr | Fabp1 | Tat | Apoh | Apoc3 | | | | | |
| Endothelial cell | Vwf | | | | | | | | | | |
| Cholangiocyte | Bicc1 | Glis3 | Ddit4l | Slc5a1 | Krt7 | Sox9 | Krt9 | | | | |
| Macrophage | Spp1 | Cd63 | Lyz1 | Ccr2 | Fcgr1 | Itgam | | | | | |
| Pericentral hepatocyte | Pck1 | Ttr | Fabp1 | Tat | Glul | Cyp2e1 | Oat | Slc1a2 | | | |
| Monocyte | Chil3 | F13a1 | Cxcr3 | Klra2 | Gm9733 | | | | | | |
| Fibroblast | Dpt | Itgbl1 | Svep1 | Gsn | Mgp | Eln | Cd34 | Mfap4 | Entpd2 | Fbln2 | Col15a1 |
| Plasmacytoid dendritic cell | Siglech | Pgam2 | Klk1 | | | | | | | | |
| Kupffer cell | Cd5l | Clec4f | Vsig4 | Kcna2 | | | | | | | |
| Myofibroblast cell | Des | Myh11 | Actg2 | | | | | | | | |
| Endothelial cell of lymphatic vessel | Ccl21a | Mmrn1 | Meox1 | Nts | Stmn2 | Thy1 | Fgl2 | Tll1 | Fxyd6 | | |
| Periportal LSEC | Lyve1 | Stab1 | Kit | Flt4 | Efnb2 | Dll4 | Esm1 | | | | |
| Pericentral LSEC | Lyve1 | Stab1 | Kit | Flt4 | Rspo3 | Wnt2 | Thbd | Cdh13 | | | |
| Midzonal LSEC | Lyve1 | Stab1 | Kit | Flt4 | Stab1 | Stab2 | Lyve1 | Mrc1 | | | |
| Prroliferating cell | Mki67 | Pcna | Stmn1 | | | | | | | | |
| Central vein and capillary macrophage | Cd207 | Cx3cr1 | Olfml3 | Mmp13 | | | | | | | |
| T-helper 1 cell | Il4 | Zbtb16 | Mmp9 | Gm47258 | Cxcr6 | | | | | | |

| | | | | | | | | | | | | |
|---|---|---|---|---|---|---|---|---|---|---|---|---|
| central vein endothelial cell | Rspo3 | Vwf | Selp | | | | | | | | | |
| myeloid dendritic cell | Xcr1 | Itgae | Naaa | Flt3 | | | | | | | | |
| LSEC/endothelial cell of hepatic sinusoid | Lyve1 | Gatm | Stab2 | Ptprc | Kit | Hgf | | | | | | |
| Mature NK T cell | Klra3 | Klra8 | Klra4 | Eomes | | | | | | | | |
| B T cell doublet | Fcer2a | Cd79a | Pax5 | Ms4a14 | Fcmr | Ebf1 | Bank1 | Cd3e | Cd3d | Themis | Cd6 | Trac |
| B cell | Fcer2a | Cd79a | Pax5 | Ms4a14 | Fcmr | Ebf1 | Bank1 | | | | | |
| T-helper 17 cell | Ramp3 | Ltb4r1 | Actn2 | Tmem176a | Icos | Il17r | | | | | | |
| Natural killer cell | Klra3 | Klra8 | Klra4 | Eomes | | | | | | | | |
| Naive T cell | Sell | Ccr7 | Tcf7 | Lef1 | Nell2 | Pask | Flt3lg | Cmak4 | Mal | | | |
| Portal vein endothelial cell | Gja5 | Adgrg6 | Ly6c1 | Pdlim3 | | | | | | | | |
| T cell | Cd3e | Cd3d | Themis | Cd6 | Trac | | | | | | | |
| Hematopoietic cell | Ptprc | Coro1a | Rac2 | Cd53 | Laptm5 | Cxcr4 | Lcp1 | | | | | |

**Table S4: Different levels of annotation.** The left side is used for Supplementary Figure 1 and the right side for deconvolution.

| celltype_pub_KC | celltype_merged |
|---|---|
| B T cell doublet | B T cell doublet |
| B cell | B cell |
| Kupffer cell | Kupffer cell |
| LSEC | periportal LSEC, pericentral LSEC, midzonal LSEC, endothelial cell of hepatic sinusoid |
| T cell | T-helper 1 cell, mature NK T cell, T-helper 17 cell, naive T cell, T cell |
| cholangiocyte | cholangiocyte |
| endothelial cell | endothelial cell, endothelial cell of lymphatic vessel, central vein endothelial cell, portal vein endothelial cell |
| fibroblast | fibroblast, myofibroblast cell |
| hematopoietic cell | hematopoietic cell |
| hepatocyte | periportal hepatocyte, hepatocyte, pericentral hepatocyte |
| myeloid leukocyte | macrophage, monocyte, central vein and capillary macrophage, myeloid dendritic cell |
| natural killer cell | natural killer cell |
| not specified | mixed, proliferating cell |
| Plasmacytoid dendritic cell | plasmacytoid dendritic cell |
